# A systematic review and comparison of automated tools for quantification of fibrous networks

**DOI:** 10.1101/2022.09.08.507154

**Authors:** Judith J. de Vries, Daphne M. Laan, Felix Frey, Gijsje H. Koenderink, Moniek P.M. de Maat

## Abstract

Fibrous networks are essential structural components of biological and engineered materials. Accordingly, many approaches have been developed to quantify their structural properties, which define their material properties. However, a comprehensive overview and comparison of methods is lacking. Therefore, we systematically searched for automated tools quantifying network characteristics in confocal, stimulated emission depletion (STED) or scanning electron microscopy (SEM) images and compared these tools by applying them to fibrin, a prototypical fibrous network in thrombi. Structural properties of fibrin such as fiber diameter and alignment are clinically relevant, since they influence the risk of thrombosis. Based on a systematic comparison of the automated tools with each other, manual measurements, and simulated networks, we provide guidance to choose appropriate tools for fibrous network quantification depending on imaging modality and structural parameter. These tools are often able to reliably measure relative changes in network characteristics, but absolute numbers should be interpreted with care.

## Introduction

Fibrous networks are critical structural components of many biological and engineered materials and are therefore studied in a wide range of fields. In a biomedical context, important examples include collagen fibers found in the extracellular matrix of mammalian connective tissues^1^, fibrin fibers found in thrombi^2^, and neurons in the brain^3^. Material scientists widely use fibrous networks to engineer hydrogels^4^, paper, and textiles^5^. The structural characteristics of fibrous networks, such as the fiber length, diameter, density and alignment, dictate the physical properties of these materials, including their elastic modulus, strength, and permeability^6^. The structure of fibrous networks is commonly determined by imaging, in particular by scanning electron microscopy (SEM) or confocal microscopy. SEM provides high resolution, but requires extensive sample preparation such as dehydration and sputter coating, which can affect network properties and introduce imaging artefacts^7^. Preparation of samples for confocal microscopy is less invasive, but this method has a limited resolution and can therefore not reliably measure the diameter of fibers, which are often below 200 nm in biological materials^8^ and synthetic hydrogels^9^. Recently developed super-resolution methods such as stimulated emission depletion (STED) microscopy offer a good alternative, but are so far not exploited much for fibrous networks. STED microscopy selectively switches off fluorophores around the focal point, thereby increasing resolution below 50 nm and improving the differentiation of separate fibers^10^. An advantage of STED microscopy over other super resolution techniques is that it is applicable to samples prepared in the same way as for confocal microscopy.

Many different approaches have been developed for the (semi-)automated quantification of fibrous network characteristics from microscopy images, though often in a specific and narrow context, such as collagen or (synthetic) nanofibers^11^ and neurites in neurons^12^. It is unknown to what extent these tools can be used across fields for other types of fibrous networks. More generally, it is unclear which methods are suitable for which structural characteristics because a comprehensive overview is missing. Therefore, our first aim is to provide an overview of publicly available automated tools that can be used to quantify fibrous network characteristics. Next, we systematically tested these tools on confocal, STED, and SEM images of fibrin networks, a representative example of a biological fibrous network with high clinical relevance. Fibrin is the main structural component of the thrombus that forms upon blood clotting. Structural properties of fibrin are important determinants of the disease burden and mortality associated with various diseases, such as cardiovascular disease and inflammatory diseases such as COVID-19^13–15^. The fibrin network forms after activation of the coagulation cascade, ultimately leading to the cleavage of fibrinogen molecules into fibrin monomers that laterally and longitudinally associate into thick fibers that form branched networks^16^. Characteristics of the fibrin network, such as fiber thickness, pore size, and number of branch points, determine among others the risk of embolization and the susceptibility of thrombi to fibrinolysis^17^. For instance, in patients with thrombotic disease, more compact thrombi are observed that are characterized by many thin fibers, a dense fibrin network with small pores, and a large number of branch points^18^. This results in decreased permeability for proteins of the fibrinolytic system, enhancing resistance to breakdown of the thrombus^19^.

Here we systematically searched the literature for available automated tools that can be used to quantify characteristics of fibrous networks. By testing these tools on confocal, STED, and SEM images of fibrin networks, we provide guidance to choose appropriate tools for the quantification of fibrous network characteristics.

## Results

### Study selection

A total of 9212 articles was found in the literature search (see Supplementary Information for the full search terms). After deduplication, 5799 articles were screened based on title and abstract (Figure 1). From the 1055 articles read in full text, we included 144 articles in which an automated tool was used that was publicly available as a script, program, or plugin. In addition, while reading these articles full text, we identified 39 other relevant articles in the references which were also included. In total, we identified 75 different automated tools or scripts in 183 papers.

**Figure 1.**
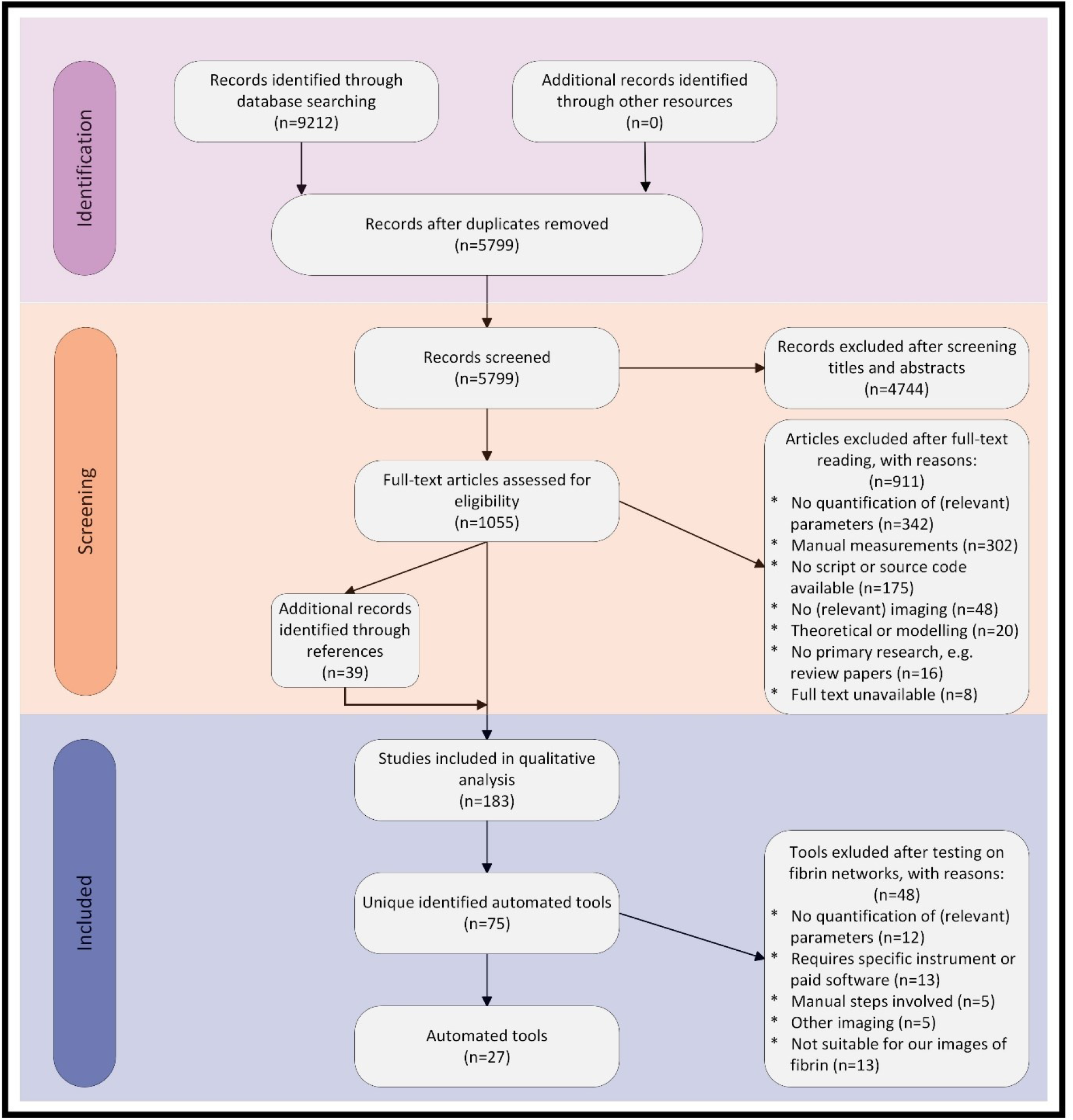
PRISMA (Preferred Reporting Items for Systematic Reviews and Meta-Analyses^70^) flow diagram of study selection. In the identification phase, we systematically searched the literature to identify articles using automated tools that quantify fibrous network characteristics. In the screening phase, articles were first screened based on title and abstract. Next, articles were read full-text to assess their eligibility. Finally, we included 183 articles in which automated analysis tools were used, of which we extracted 75 different tools. We applied these tools to fibrin, a prototypical fibrous network example present in thrombi. Of the 75 tools, 27 could be used on our confocal, stimulated emission depletion (STED), or scanning electron microscopy (SEM) images. The results of these tools were compared to each other, manual measurements, and simulated networks.

### Testing of the automated tools

The selected tools were tested on our confocal, STED, and SEM images of fibrin networks to check whether they can be used on fibrin networks imaged using one of these imaging techniques. From the 75 tools, 48 tools were excluded. Reasons for exclusion during testing were, among others, that some tools required manual processing steps, could only be used on images from specific devices, needed paid software, or were not suitable for our images of fibrin fibers (Supplementary Table I). This last category includes tools that for example looked for cross-sections of fibers^20^, needed multi-channel images^21,22^, or needed specific kinds of structures, such as cell bodies in neurons^12^.

### Comparison of the automated tools

Next, the automated tools that could be used on our fibrin network images were applied to 100 confocal (50 of fibrin networks formed under static conditions and 50 of fibrin networks formed under flow), 50 SEM, and 50 STED images of fibrin networks (see Supplementary Figure I-IV for visual presentations of the applied tools). Many of the automated image analysis tools were originally developed for collagen or nanofibers (Table 1). Most tools were suitable for confocal, STED, and SEM images and could be used with a batch processing option. In addition, most tools quantified network characteristics in 2D, while some were able to also, or solely, use 3D image stacks. Most of the identified tools were able to quantify fibrin fiber diameter, fiber length, or alignment (Table 2). Three of the 27 tools were not included in the comparison analyses. FIRE^23^ was not used, since CT-FIRE^24^ was used instead, which is an extension of FIRE. Qiber3D^25^ and the 3D directional variance algorithm^26^ could only be used on 3D image stacks. However, all other tools were applied to 2D images, so we did not include Qiber3D and the 3D directional variance algorithm in the comparison analyses.

**Table 1.**
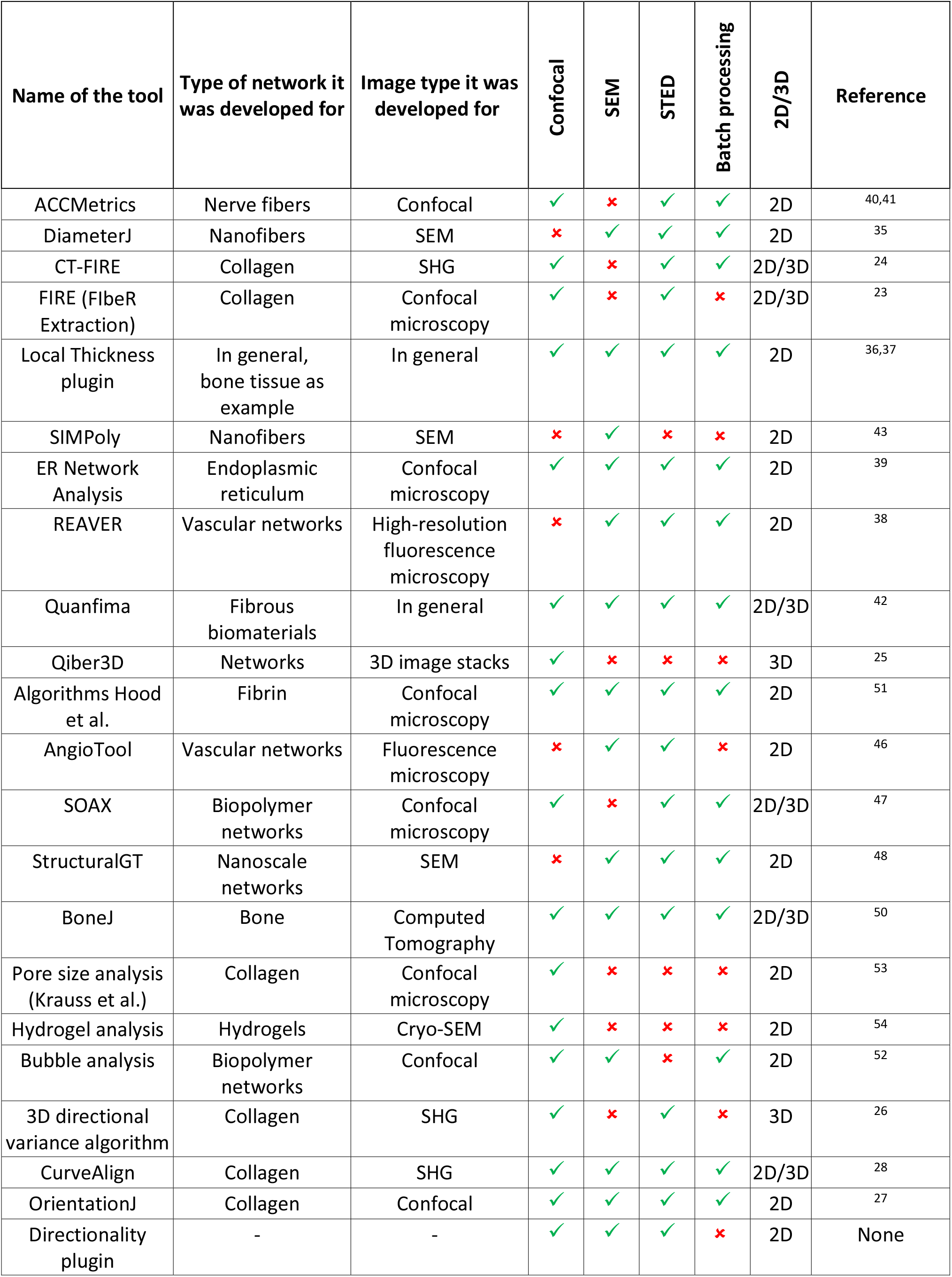
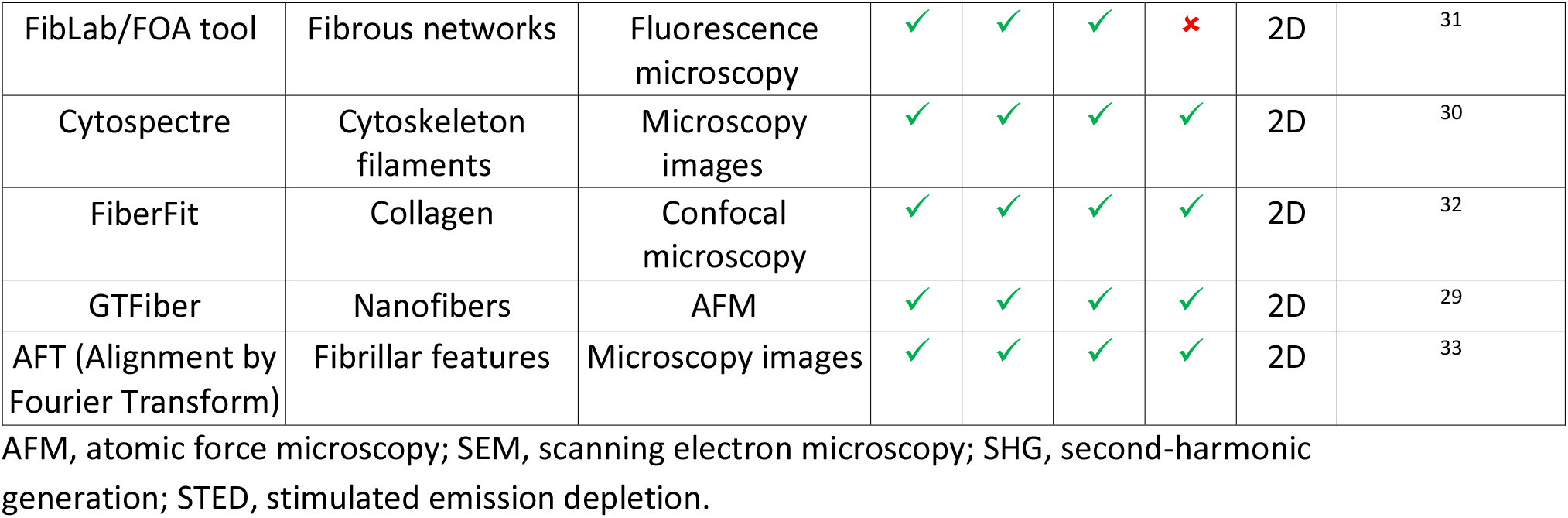
Overview of identified automated tools that can be used to quantify fibrin fiber network characteristics.

**Table 2.**
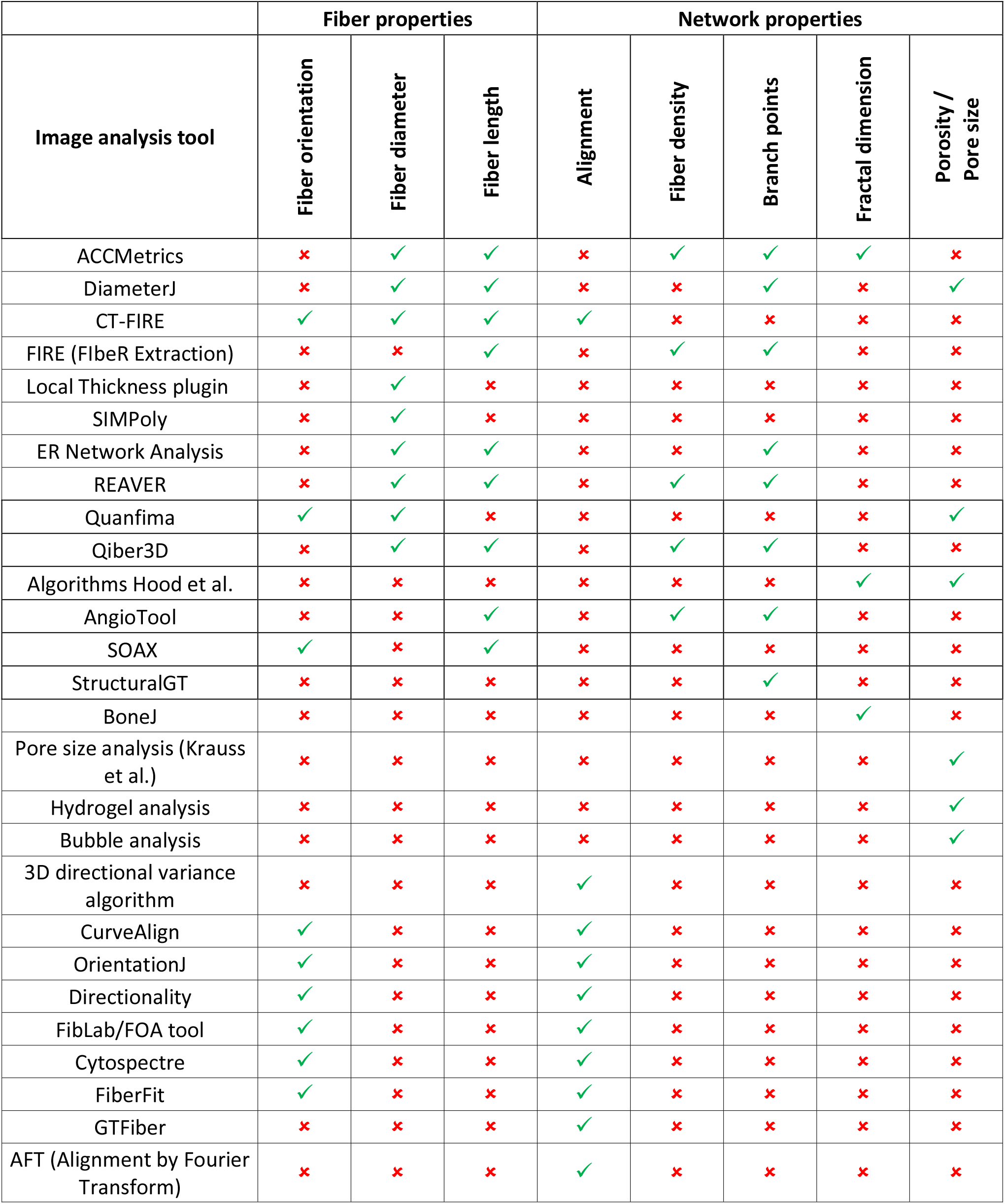
Overview of quantified characteristics per tool.

### Fiber alignment

We tested nine tools that could quantify the alignment of the fibrin network (Supplementary Table II): OrientationJ^27^, CurveAlign^28^, GTFiber^29^, CT-FIRE^24^, Cytospectre^30^, Directionality, FOA tool^31^, FiberFit^32^, and Alignment by Fourier Transform (AFT)^33^. Most of the methods measured global alignment (within the whole image), while OrientationJ, GTFiber, and AFT used local alignment measurements (within the local neighborhood). To compare these tools, we applied them to confocal images of fibrin fibers aligned by applying shear flow with different shear rates. Visual examples of the applied tools are shown in Supplementary Figure IV. Supplementary Figure V shows the 50 images with their corresponding nematic order parameter (S) as calculated by OrientationJ. OrientationJ was used as reference because of its proven sensitivity for fiber alignment^34^.

OrientationJ and the CurveAlign, GTFiber, CT-FIRE and AFT tools provide an orientation parameter that describes the alignment with a value between 0 (randomly aligned) and 1 (perfectly aligned). Strong correlations were found between these five tools, with good agreement in absolute levels between OrientationJ and CurveAlign and between CT-FIRE, GTFiber, and AFT (Supplementary Table II and Supplementary Figure VI). CT-FIRE, GTFiber, and AFT report levels that are clearly higher than OrientationJ and CurveAlign. Cytospectre also returns a value between 0 and 1, but in this case the alignment parameter is defined such that 1 means randomly aligned and 0 means perfectly aligned. This is opposite to the other tools, which explains the negative correlations. FiberFit returns the fiber dispersion parameter, which is analogous to the reciprocal of variance in fiber orientation, therefore low values mean disordered networks and large values mean aligned networks. FiberFit shows a significant correlation with OrientationJ. Finally, the directionality plugin and the FOA tool result in parameters describing the spread of the orientation distribution. Lower values represent better aligned fibers, while higher values represent more variation and therefore less aligned fibers. Directionality and the FOA tool appear to be rather insensitive to alignment as quantified by OrientationJ.

Additionally, we applied the tools to synthetic images generated based on a semicircular von Mises probability density function^32^ with known dispersion parameters (Figure 2A and Supplementary Table III). Since all tools report different outcome parameters, we could not compare absolute values to the known dispersion parameter. Similar to the results in the confocal images, we did see very strong significant correlations (r>0.85) of the results of almost all tools with the known dispersion values, except for the FOA tool (r=-0.73, p<0.05). The Directionality plugin did not provide useful results from these images, since no reliable fit could be made to the histogram.

**Figure 2.**
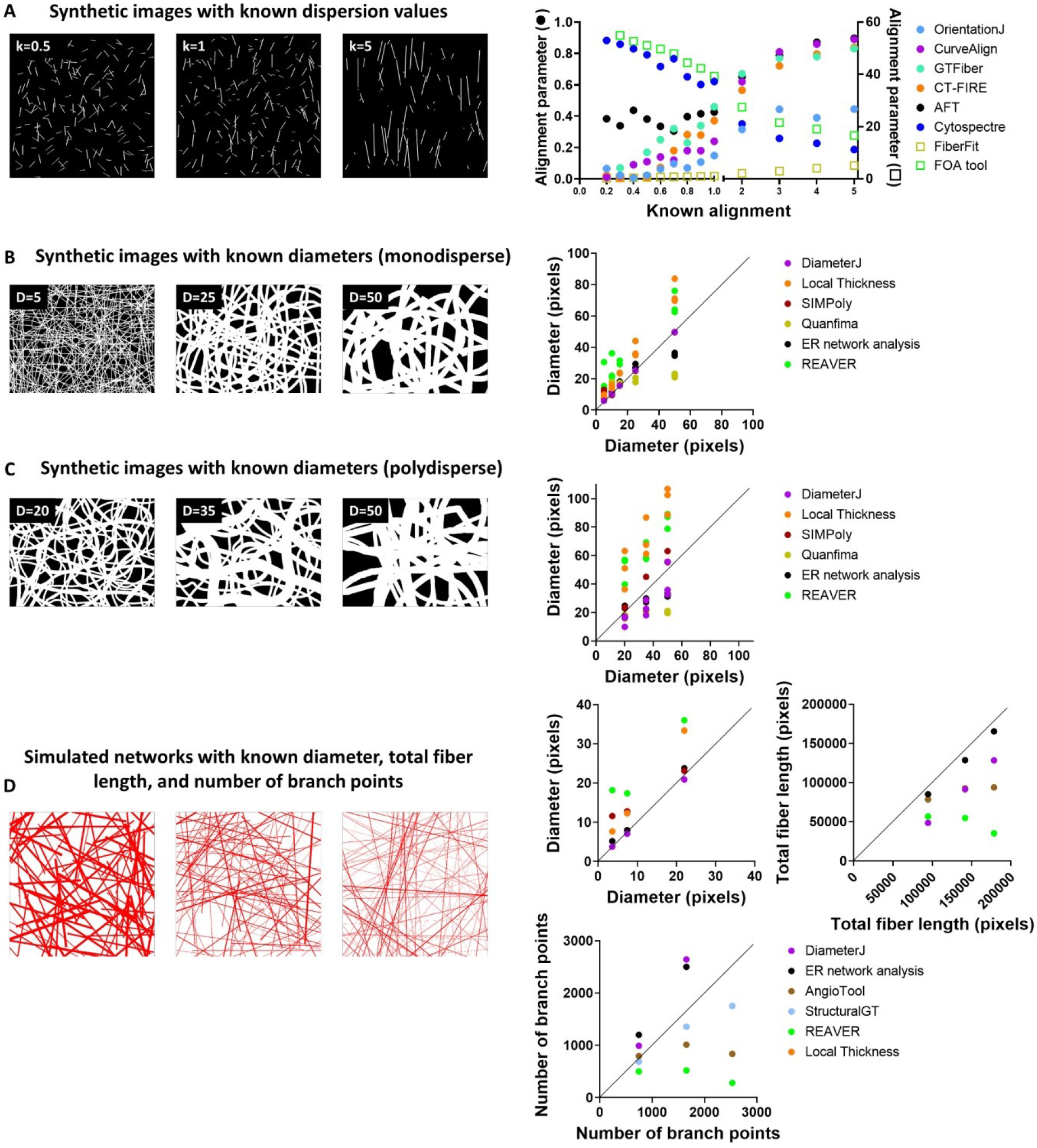
Examples of (A) synthetic images with known dispersion (k) values^32^, synthetic images with (B) monodisperse or (C) polydisperse diameters^44^, and (D) simulated networks. Correlation plots show the known values on the x-axis and the results of the different tools on the y-axis. The solid lines indicate the x=y-lines. Every dot represents the result of one image. In B and C, three different images per diameter value were used, while in A and D, only one image per value was used.

### Fiber diameter

To quantify fibrin fiber diameter, we identified eight automated tools: DiameterJ^35^, Local Thickness plugin^36,37^, REAVER^38^, ER network analysis^39^, ACCMetrics^40,41^, CT-FIRE^24^, Quanfima^42^ and SIMPoly^43^. Because a high image resolution is needed to quantify fibrin fiber diameters, we only tested these tools on STED and SEM images (Figure 3A). For SEM images, we benchmarked the results from the automated tools against manually measured fiber diameters^8^. Both in SEM and STED images, almost all tools showed strong correlations with each other (Figure 3B and Supplementary Table IV and V). SIMPoly and the ER network analysis found similar absolute values as the manual measurements in SEM images. By contrast, DiameterJ systematically underestimated fiber thickness. The mean diameter measured by DiameterJ was 17% lower than the manual measurement. The Local Thickness plugin and REAVER showed no or weak correlations with the manual measurements, as well as a large overestimation (31%) of the fiber thickness in the case of REAVER. In the STED images, we used the results of the ER network analysis for reference since the mean value of the diameter calculated by this tool was closest to the manual measurement of the diameter in the SEM images. All other tools showed strong correlations with the ER network analysis tool and with each other in STED images. The absolute diameter values in STED images ranged from 200 nm (measured by CT-FIRE) to 400 nm (measured by REAVER).

**Figure 3.**
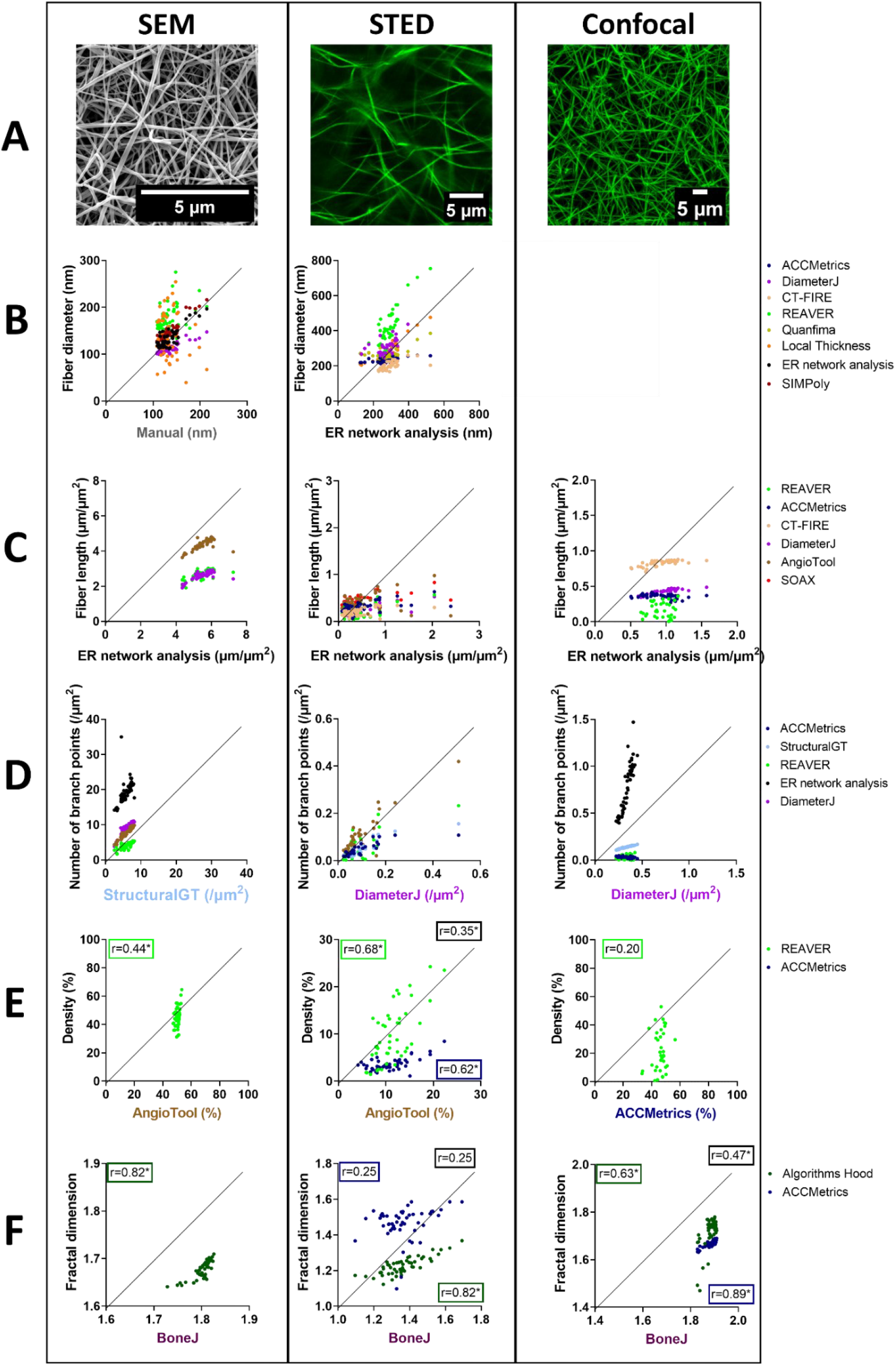
Correlation between the results of the different tools applied to SEM, STED, and confocal images (A) to quantify fibrin fiber diameter (B), fiber length (C), number of branch points (D), fibrin network density (E), and fractal dimension (F). The solid lines in (B-F) indicate the x=y-lines. B) Manual measurements of fiber diameter were used as reference on the x-axis in the SEM images, while the fiber diameters quantified by the ER network analysis were used as reference in the STED images, since its absolute values were closest to the manual measurements in SEM. Every dot presents the mean fiber diameter per image. C) For fiber length measurements, the results of the ER network analysis were used as reference, since the results from this tool were closest to known values in synthetic images. Total fiber length measured by the different tools was divided by the surface area of the image, to be able to compare absolute values between different imaging methods. Every dot represents the fiber length per μm^2^ in one image. D) For the quantification of the number of branch points, the numbers calculated by StructuralGT (SEM) and DiameterJ (STED and confocal) were used as reference, since these tools performed best on simulated networks with known number of branch points. The total number of branch points was divided by the surface area of the image. Therefore, every dot represents the number of branch points per μm^2^ in one image. E, F) Fibrin network density and fractal dimension were quantified by only two or three tools. Therefore, Pearson’s correlation coefficients (r) are given for every combination of tools in the graph, indicated by boxes in the color of the dots of the correlation of that specific tool with the tool on the x-axis, or in black boxes when two tools on the y-axis are correlated; *p<0.05.

To further check the performance of the tools, we next tested them also on synthetic images of fibrous networks drawn using the freehand tool in Inkscape^44^ with either known and validated monodisperse fiber diameters between 5 and 50 pixels (Figure 2B) or polydisperse fiber diameters with an average of 20, 35, or 50 pixels (Figure 2C), and to simulated networks with known characteristics (Figure 2D and Supplementary Figure VII), generated using an algorithm in Python (see Supplementary Information for the Python code). We used both synthetic images of fibers that were published and validated previously for DiameterJ and FiberFit, as well as fibrous networks that we simulated ourselves to be able to perform unbiased comparisons. DiameterJ performed well in synthetic images and the simulated networks with constant fiber diameters, while diameter values were underestimated (on average by 44%) in the synthetic images with polydisperse fiber diameters (Supplementary Table VI). This limitation is a consequence of the intersection subtraction of DiameterJ^35^. Thicker fibers are more likely to have more intersections due to their increased size. Therefore, thicker fibers will have less influence on the mean diameter than thinner fibers, which results in an underestimation of the mean fiber diameter. Fibrin networks are usually polydisperse in terms of fiber diameters^45^, making DiameterJ less suitable for absolute diameter quantification. The Local Thickness plugin and REAVER were sensitive to changes in fiber diameters in the synthetic images, but significantly overestimated (mean) fiber thickness by respectively 79% and 121%. SIMPoly performed well in images containing fibers with a thickness between 10 and 50 pixels, while the fiber thickness in the images with the thinnest fibers was not correctly measured. This is probably caused by an unsuccessful detection of the thinner fibers. The ER network analysis on the other hand showed correct values for fibers with a diameter up to 25 pixels, while fibers of 50 pixels were underestimated. Visual inspection showed that in very thick fibers, sometimes two centerlines were drawn, which might explain this underestimation. Quanfima was rather insensitive to changes in fiber diameter. Overall, we recommend to use ER network analysis or SIMPoly to measure fibrin fiber diameters.

### Fiber length

We found seven different tools that quantify fiber length in confocal, STED, or SEM images: DiameterJ^35^, AngioTool^46^, REAVER^38^, ER network analysis^39^, CT-FIRE^24^, ACCMetrics^40,41^, and SOAX^47^ (Figure 3C). The results of these tools showed strong correlations with each other when applied to STED and SEM images, but not for confocal images (Supplementary Tables VII-IX). Absolute values of fiber length, normalized against the surface area of the image, were higher in SEM images compared to STED and confocal images, likely since SEM images show fibers from multiple layers whereas STED and confocal show fibers only within the confocal slice. Applying the automated tools to simulated networks showed that the ER network analysis returned values very similar to the known values (Figure 2D and Supplementary Table X). DiameterJ was sensitive to changes, but underestimated fiber length, while AngioTool and REAVER were not sensitive to fiber length. We hence recommend to use the ER network analysis to automatically determine fiber length.

### Number of branch points

To automatically quantify the number of branch points, we found six tools: DiameterJ^35^, AngioTool^46^, StructuralGT^48^, REAVER^38^, ER network analysis^39^, and ACCMetrics^40,41^. Again, strong correlations were found between the results of the tools in both SEM and STED images, with remarkably higher absolute numbers of branch points per μm^2^ in SEM than in STED or confocal images (Figure 3D and Supplementary Tables XI-XIII). In confocal images, significant associations were present between DiameterJ, StructuralGT, and the ER network analysis. However, absolute numbers of the different tools showed a very broad range, suggesting that confocal imaging is not a reliable tool for measuring branch points. In SEM images, REAVER reported a significantly lower number of branch points compared to the majority of tools, while the ER network analysis resulted in a significantly higher number. In STED images, also a large range of numbers was found: between 30 (StructuralGT and ACCMetrics) and 70 (DiameterJ and AngioTool). Upon inspection of the fiber tracing results, we observed that StructuralGT and ACCMetrics did not detect all fibers and therefore all branch points present in STED images, while DiameterJ and AngioTool performed better (Supplementary Figure III). When analyzing the simulated networks, we observed the best results for StructuralGT (Figure 2D and Supplementary Table XIV). DiameterJ and the ER network analysis overestimated the number of branch points, while AngioTool and REAVER were insensitive to changes in the number of branch points. Based on visual inspection and the results from the simulated images, we recommend to use StructuralGT in SEM images and DiameterJ in STED images to automatically measure the number of branch points.

### Fiber density

We found three different tools to quantify the density of the fibrin network: AngioTool^46^, REAVER^38^, and ACCMetrics^40,41^. Density is defined as the fraction or percentage of the area occupied by fibers. Similar to the branching analysis, significant correlations were observed between the quantifications in SEM and STED images, while results in confocal images did not significantly correlate (Figure 3E). Absolute numbers strongly varied between different automated tools with no tool clearly being most reliable (Supplementary Table XV).

### Fractal dimension

Fractal dimension can be used to investigate complexity of the network. It measures how details in an object change with the scale at which it is measured^49^. BoneJ^50^, algorithms developed by Hood et al.^51^, and ACCMetrics^40,41^ were able to analyse the fractal dimension in our fibrin networks. In SEM and STED images, a strong correlation between the fractal dimension calculated by BoneJ and Algorithms of Hood et al. was found, while ACCMetrics gave different results in STED images (Figure 3F and Supplementary Table XVI). In confocal images, the three tools correlated strongly.

### Pore size and porosity

Finally, we found some tools that automatically measured pore size or porosity, the size or fraction of pores in the network. In SEM images, we could only use DiameterJ^35^ and the Bubble analysis tool^52^ to measure mean pore size. DiameterJ reports mean pore area in μm^2^, while the Bubble analysis tool reports the mean pore diameter in μm. No correlation between both measurements was found (r= −0.12, p=0.40) (Supplementary Figure VIIIA). In confocal images, six different tools were found which report pore size, pore diameter, or porosity: DiameterJ^35^ (mean pore area and porosity), Pore size analysis by Krauss et al.^53^ (mean pore diameter), Bubble analysis^52^ (mean pore diameter), Hydrogel pore size analysis^54^ (mean pore area, mean pore diameter, and porosity), Algorithms by Hood et al.^51^ (porosity), and Quanfima^42^ (porosity) (Supplementary Table XVII and Supplementary Figure VIIIB-C). Except for the Hydrogel analysis, the porosity measurements correlated quite well between the different tools. Pore size measurements by the different tools showed only significant correlations when the same parameter was calculated, either diameter or pore area. The pore diameters measured by the Pore size analysis by Krauss et al.^53^ and the Bubble analysis^52^ did also not correlate with the porosity measurements, while the tools calculating pore area^35,54^ did show significant correlations with the porosity measurements.

## Discussion

We systematically searched the literature for all available tools that automatically quantify structural characteristics of fibrous networks from either electron or light microscopy images. Next, we compared the results from the identified automated tools when applied to confocal, STED, and SEM images of fibrin networks, a prototypical fiber with biological and clinical relevance. We found 27 different automated tools that were publicly available and could be applied to our images. Of these tools, a minority were originally developed for fibrin (Table 1). This shows the importance of making research tools available and findable across different fields.

Using the identified tools, we measured fiber alignment, fiber diameter, fiber length, number of branch points, fiber density, fractal dimension, pore size, and porosity. It was striking that the results of the different automated tools often showed good correlations, while the absolute numbers varied a lot. This suggests that most tools are able to measure relative changes in fibrous network characteristics but absolute numbers should be interpreted with care. Therefore, we recommend that comparison of fibrous network characteristics should always be performed using the same type of microscopy image and analysis tool. Another observation we made is that the correlations between results from different automated tools were often stronger in the images with high resolution (SEM and STED) than in the confocal images. For confocal images, the optical diffraction limit, being comparable to the size of the fibers, probably limits the reliability of automated analysis of fiber and junction features. Moreover, out of focus light might also influence the measurements. However, confocal images may be preferred in measurements of fiber alignment, network density and fractal dimension because of their larger field of view compared to electron microscopy. Furthermore, absolute numbers of fiber length per μm^2^, number of branch points per μm^2^, and density were higher in SEM than in STED and confocal images. This is likely caused by the fact that the scanning electron microscope images the surface of fibrous networks and captures fibers from multiple layers, whereas STED and confocal microscopy optically filter out fibers from above and below the confocal plane. The smaller fiber diameter in SEM than in STED images can probably be explained by shrinkage of the clot in SEM due to the preparation, or some remaining out of focus light in STED images. Finally, our systematic search revealed that many articles use automated ways of measuring fibrous network characteristics, without making their tools or scripts available. Therefore, to increase standardization and transparency, open access of newly developed tools should be promoted.

Most of the identified tools were developed for the quantification of fiber alignment, fiber diameter or fiber length of fibrous networks. Alignment of fibers in fibrous networks affects cellular behavior, such as cell attachment, migration, and differentiation, which is important in tissue engineered scaffolds^55^, wound healing, and - in case of fibrin - the susceptibility of thrombi to fibrinolysis^56,57^. We tested nine different tools to automatically quantify alignment of fibrin fibers. These tools either resulted in a nematic order parameter between 0 and 1 or a parameter quantifying the spread of the orientation distribution. In general, the different tools showed good correlations, showing that most tools can be reliably used to measure fibrin fiber alignment in confocal images. Using synthetic images with known fiber dispersion confirmed that most of the tools, except for the Directionality plugin, performed well in the quantification of fiber alignment. We recommend a tool that returns a nematic order parameter, since this provides a sensitive measure between 0 and 1 that can be linked directly to theoretical models that connect fiber alignment to mechanical properties^34,58^. When also taking into account the ease of use of the tools and the possibility to adapt parameters, our preferences to quantify fiber alignment in fibrin networks go towards OrientationJ^27,59^ (ImageJ plugin), CurveAlign^28^ (MATLAB-based standalone application), or FiberFit^32^ (Python-based standalone application).

Until recently, quantification of fiber diameters was often done manually, which is inefficient and sensitive to observer bias^8,60^. We tested eight available tools to automatically measure fiber diameter. Our analyses showed that both in SEM and STED images, fiber diameter quantifications were strongly correlated among different methods, while absolute values differed markedly. Using manual measurements and synthetic and simulated images, we showed that SIMPoly and the ER network analysis tools reported values closest to the true diameters. Therefore, we recommend to use SIMPoly in SEM images and the ER network analysis in SEM or STED images. However, both tools showed some variation in performance based on the width and polydispersity of the fibers. The optimal thickness of fibers in images is between 10 and 25 pixels for both tools, which would require a pixel size between 10 and 25 nm for typical fibrin fibers. SIMPoly is a MATLAB-based tool that is very user-friendly^43^. The ER network analysis tool provides more user control because it has many options that can be adjusted, but it therefore requires more testing by the user for their specific images^39^.

Fiber length, the number of branch points, and fiber density are other characteristics which are often used to describe the organization of fibrous networks. Increased branching is associated with thinner fibers, which is characteristic for more dense fibrin networks^61^. Our analyses showed that in SEM and STED, good correlations were found between the different tools. For fiber length, best results were obtained with the ER network analysis, while StructuralGT gave the best results in the quantification of the number of branch points. Our results indicate that electron microscopy is more suited than confocal imaging for precise quantification of fiber length and the number of branch points. For fiber density, absolute numbers strongly varied between the three different automated tools with no tool clearly being most reliable.

The fractal dimension of fibrin networks is known to increase with increasing thrombin concentration and a denser fibrin network structure^62–64^. Moreover, a recent study suggests that fractal dimension of the fibrin network is a biomarker for a high risk of venous thrombosis^64^. In our analysis, we found only three tools that measured fractal dimension, with variable results. More research should be performed to determine whether this characteristic can be reliably measured in microscopy images to describe network complexity of fibrous networks, for example by comparing it to fractal dimension quantifications using rheology^65^.

Finally, pore size and porosity were quantified by six different tools. When measuring pore size, only moderate correlations were found between the results from the different tools. In contrast, porosity measurements of different tools were more strongly correlated. Correlations between pore size and porosity measurements were best when the pore area was calculated instead of the pore diameter, suggesting a circle might not be the best description of a pore. A major drawback of the included tools is the fact that the pores are mostly measured in 2D images, while it is important to measure pores in 3D to get reliable information since pores can have different sizes in all three dimensions. Pore size and porosity as measured by DiameterJ were indeed shown to be poorly correlated with functional permeability measurements of plasma clots^8^. While pore sizes have been measured in 3D^66–68^, and it was shown that it is possible to work with 2D images while correcting for the missing third dimension^69^, these tools are unfortunately not publicly available. We conclude that automated tools exist that can measure porosity and/or pore size in 2D images, preferably by measuring pore area, but that better tools should become publicly available.

In conclusion, we have made an overview of a large number of automated quantification tools for fibrous networks and systematically compared them on fibrin networks, a prototypical example of a fibrous network in thrombi. We conclude that fiber diameter can be measured reliably and efficiently by SIMPoly or the ER network analysis in SEM or STED images (Figure 4). For fiber alignment, we recommend using OrientationJ, CurveAlign, or FiberFit on confocal images. Fiber length can be measured using the ER network analysis, while we recommend DiameterJ or StructuralGT to measure the number of branch points in SEM or STED images. For the other network characteristics, such as fractal dimension or pore size, more development and open access publication of automated tools is needed. We anticipate that our analysis provides researchers useful guidelines in their search for suitable automated analysis tools of fibrous networks in different fields, from thrombosis and hemostasis to cancer research, regenerative medicine, materials science, and neuroscience.

**Figure 4.**
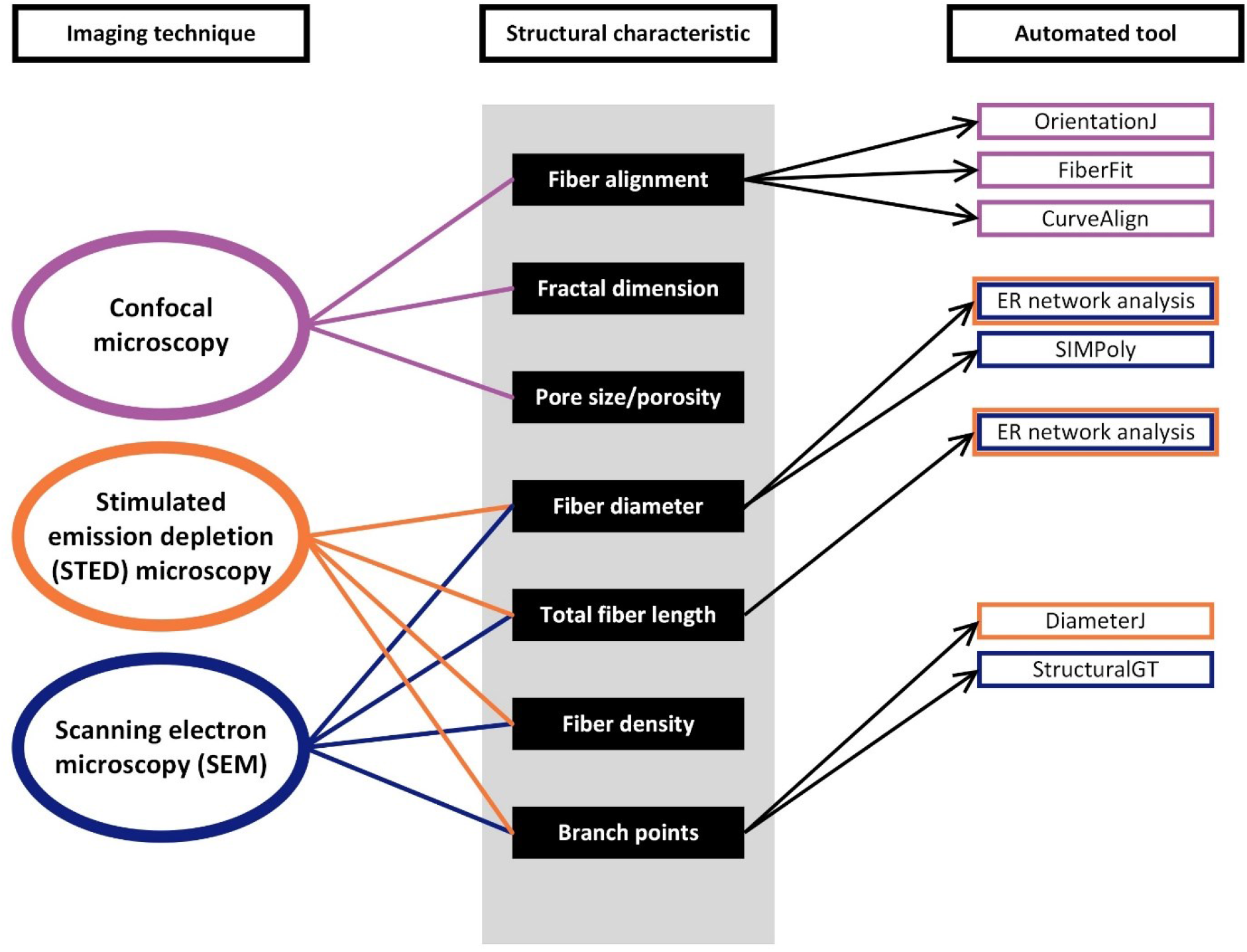
Overview of fibrous network characteristics with their appropriate imaging technique and recommended tool for automated analysis. For every network characteristic (middle panel), we indicated the recommended imaging technique (left panel) by connecting the imaging technique and characteristic using colored lines. Also, we recommend automated tools by connecting the network characteristic to an automated tool (right panel). The color of the boxes in the right panel correspond to the suitable imaging technique, e.g. the ER network analysis can be used for both STED and SEM images, while StructuralGT is only recommended for SEM images. For fractal dimension, pore size/porosity, and fiber density, we could not determine the best tool, so no recommendation is given.

## Online Methods

The systematic review was performed according to the Preferred Reporting Items for Systematic Reviews and Meta-Analyses (PRISMA) guidelines^70^.

### Article Search

We conducted a systematic literature search in the Embase, Medline-Ovid, Cochrane Library, and Web of Science databases on April 2nd, 2021; and the search was repeated on November 7, 2021 and May 10, 2022. The search strategy included different types of fibrous networks, such as fibrin, collagen, and nanofibers, in combination with image analysis terms (e.g. algorithm or automated analysis), different microscopes (e.g. fluorescence, confocal, scanning electron microscopy), and network characteristics (e.g. diameter, pore size, orientation, length). For the full search terms, see the Supplementary Information.

### Study Selection

After deduplication, the search resulted in 5799 results. Two researchers (J.J. de Vries and D.M. Laan) independently screened these articles. In the first step, articles were included based on title and abstract. Subsequently, these abstracts were read full text and articles that did not match the research questions were excluded. Reasons for exclusions were: manual measurements, no quantification of (relevant) parameters, no (relevant) imaging, or when the paper was only theoretical. In addition, reviews and articles not available in full text were excluded. While reading the articles, additional relevant articles based on the bibliography were also assessed for inclusion. In case of disagreement between the two researchers, consensus was reached through discussion.

### Data Extraction

To prevent bias, data were independently extracted from the articles by two researchers (J.J. de Vries and D.M. Laan). The following data was collected: first author, publication year, name of the used tool, type of network the tool was used on, imaging modality used to make the images, and quantified characteristics (alignment, fiber diameter, fiber length, fiber density, number of branch points or junctions, pore size and/or porosity).

### Testing and comparison of the automated tools

After identification of the automated tools, these tools were tested on different sets of images of fibrin networks to assess whether the tools worked on these images and resulted in automated quantification of relevant characteristics. Tools were excluded when manual steps such as manual tracing were needed, when only images from specific devices could be used, when the software was commercial, or when specific requirements were needed that were not present in our fibrin network images (e.g. multiple imaging channels or specific shapes, such as a neuron body). Next, tools that worked on our fibrin images were applied on a test set of fibrin network images and results of the different tools were compared. For each quantified characteristic, we assessed the correlation between results obtained using the different tools, reporting the Pearson’s correlation coefficient. Before analyzing the images using the automated tools, STED and confocal images were preprocessed by subtracting the background. A rolling ball radius of 50 pixels for STED images and 25 pixels for confocal images was used. These settings were based on the size of the largest objects in the images. In addition, other relevant preprocessing steps were used when this was deemed necessary for a good performance of the analysis tool (Supplementary Table XVIII). Most of the settings in each tool were left as their default settings, except when stated otherwise in Supplementary Table XVIII.

The first data set was a series of scanning electron microscopy (SEM) images of clots prepared from plasma from healthy controls or patients with severe postpartum hemorrhage (PPH)^8^. This study was approved by the Medical Ethics Committee of the Erasmus University Medical Center Rotterdam (MEC-2016-218) and all participants provided written informed consent. Clots were prepared and imaged as described in Daraei et al.^8^. Briefly, clots were prepared by adding 25 mM CaCl2 and 0.5 U/ml thrombin to plasma and incubating them for 30 minutes at room temperature with subsequently washing, dehydration and drying to prepare them for SEM imaging. Samples were sputter coated with a layer of gold/palladium and imaged at 10,000 times magnification at eigth randomly selected areas. The pixel size of the SEM images was 8.3 nm. For the current study, five images from six patients with severe PPH and four healthy controls were used, resulting in a total of 50 SEM images. For these images, manual measurements of fiber diameter were available, which were used to benchmark the automated analyses^8^.

The second set of images were confocal laser scanning microscopy images of clots prepared from plasma from healthy volunteers^71^ and COVID-19 ICU patients^15^. These studies were approved by the Medical Ethics Committee of the Erasmus University Medical Center Rotterdam (healthy controls: MEC-2004-251; COVID-19 ICU patients: MEC-2020-0758) and participants provided written informed consent or an opt-out procedure was in place. Clots were prepared by adding 17 mM CaCl2, 1 U/ml thrombin, and 4% fluorescently labeled fibrinogen to citrated plasma, incubating them for 1.5 hours at room temperature with subsequently washing and fixation to prepare them for confocal imaging. For each clot, at three random positions, Z-stacks were obtained using a LEICA TCS SP8 microscope (Leica Microsystems, Wetzlar, Germany). A LEICA HC PL APO CS2 40x/1.30 oil immersion objective was used. Image size was 2048×2048 pixels (138.4 μm), resulting in a pixel size of 67.6 nm. A step size of 1 μm was used for the Z-stacks, starting approximately 5 μm above the glass coverslip until approximately 20 μm above the glass. For the current study, two images of five healthy volunteers and two images of 20 COVID-19 ICU patients were used, resulting in a total of 50 confocal images.

To obtain super resolution fluorescent images, the plasma clots described in the previous paragraph were also imaged using the LEICA TCS SP8 microscope using STED settings. Approximately 10 μm above the glass, 101 images were obtained with a depletion laser at 592 nm, using a Leica HC PL APO CS2 86x/1.20 water immersion objective. Image size was 1024×1024 pixels (27.0 μm), resulting in a pixel size of 26.4 nm. For the current study, two images of five healthy volunteers and two images of 20 COVID-19 ICU patients were used, resulting in a total of 50 STED images.

The last set of images (not published) were confocal microscopy images of clots prepared in a flow chamber (Flow Chamber BV) where flow at different shear rates was used to obtain different degrees of network alignment. Control pooled plasma (VisuCon-F Frozen Unassayed Normal Control Plasma) was used, which was mixed with 17 mM CaCl2 and 1 U/ml thrombin right before perfusion through the flow chamber over a coverslip coated with 1 mg/ml fibrinogen. Z-stacks were recorded with a step size of 1 μm, starting from just above the glass coverslip using a 40x oil immersion objective (NA 1.25, HCX PL APO CS). Maximum projections of the Z-stacks and individual slices were used for fiber orientation analyses. A total of 50 maximum projections and images formed at different shear rates, ranging from 50 s^-1^ to 250 s^-1^ were used. Tools quantifying fiber alignment were only tested on this set of images.

### Comparing automated tools on synthetic images and simulated fibrous networks

To test the performance of each tool, we also applied the automated tools to digital synthetic images of branched fibrous networks with flexible fibers with known diameters, generated by Hotaling et al.^44^. We used 15 images with a monodisperse fiber diameter and nine images with polydispersity in the thickness of fibers (see Figure 2B and 2C for examples). All tools able to measure diameter were applied to these images. Additionally, we used 13 synthetic images containing rigid, short fibers with known dispersion parameters ranging from 0.2 to 5, generated by Morrill et al.^32^ (see Figure 2A for examples). These images all had a mean fiber orientation of 90°. All tools able to measure fiber alignment were applied to these images.

Finally, we used three simulated branched fibrous networks with flexible fibers with known diameters, branch point number and fiber length (Figure 2D). The algorithm to generate fibrous networks was written in Python (See Supplementary Information). For every network simulation, we set six parameters: (1) the total number of filaments “tot_number_fil”, (2) the maximum filament length “max_len”, (3) and (4) two parameters “shape” and “scale” that define the filament diameter (explained below), and (5) and (6) two parameters “geometry_x” and “geometry_y” that determine the length and height of the simulated surface, which are both set to 100 for all simulated networks. We then distributed the total number of filaments on the surface. First, we determined the filament orientation. For every filament, we drew a random angle ϕ from a uniform distribution between 0 and π with respect to the x-axis. Second, we determined the position of the filament center of mass. Therefore, for every filament, we drew the x and y coordinate of the filament center of mass from a uniform distribution between 0 and “geometry_x” and between 0 and “geometry_y”, respectively. With the filament orientation and the position of the filament center of mass, we then calculated the length of the filament within the simulated surface. Finally, we set the filament diameter, by sampling a random number from a gamma distribution that was determined by the “shape” and “scale” parameters. After we have distributed the total number of filaments on the surface, we calculated the branch points for the network by summing up all intersections between pairs of two filaments within the simulated surface. From the simulation we determined the filament length and diameter distributions and the number of branch points.

## Data availability

All confocal, STED, SEM, and simulated images can be found in a data supplement available with the online version of this article. Also the numbers obtained after applying the automated tools on the images are available in a data supplement.

## Code availability

The Python code to generate simulated images can be found in a data supplement available with the online version of this article. The software and scripts of the used methods identified in the systematic search can be found via the original publications.

## Acknowledgements

The authors wish to thank Wichor M. Bramer from the Erasmus MC Medical Library for developing and updating the search strategies, Marieke J.H.A. Kruip and Caroline S.B. Veen for providing us with the SEM images and manual measurements of the fiber diameters, Emmelien D. Hazekamp for her help with some of the MATLAB tools, and Trevor J. Lujan for providing us with test images with known dispersions.

F.F. gratefully acknowledges funding from the Kavli Synergy program of the Kavli Institute of Nanoscience Delft. G.H.K. gratefully acknowledges funding from the NWO Talent Programme which is financed by the Dutch Research Council (NWO) (project number VI.C.182.004).

## Authorship Contributions

J.J.d.V. and M.P.M.d.M. designed the study. J.J.d.V. and D.M.L. performed the literature search, performed image and data analysis, and produced the tables. J.J.d.V. produced the figures and wrote the original draft of the manuscript. F.F. developed the Python code to generate simulated images. D.M.L., F.F., G.H.K. and M.P.M.d.M. reviewed and edited the draft. G.H.K. and M.P.M.d.M. supervised the entire project. All authors read and approved the final manuscript.

## References

1. Frantz C, Stewart KM, Weaver VM. The extracellular matrix at a glance. J Cell Sci. 2010;123(Pt 24):4195–4200.

2. Mosesson MW. Fibrinogen and fibrin structure and functions. J Thromb Haemost.2005;3(8):1894–1904.

3. Ho SY, Chao CY, Huang HL, Chiu TW, Charoenkwan P, Hwang E. NeurphologyJ: an automatic neuronal morphology quantification method and its application in pharmacological discovery. BMC Bioinform. 2011;12:230.

4. Ahmed EM. Hydrogel: Preparation, characterization, and applications: A review. J Adv Res.2015;6(2):105–121.

5. Picu RC. Mechanics of random fiber networks—a review. Soft Matter. 2011;7(15):6768–6785.

6. Burla F, Mulla Y, Vos BE, Aufderhorst-Roberts A, Koenderink GH. From mechanical resilience to active material properties in biopolymer networks. Nat Rev Phys. 2019;1(4):249–263.

7. Kirk SE, Skepper JN, Donald AM. Application of environmental scanning electron microscopy to determine biological surface structure. J Microsc. 2009;233(2):205–224.

8. Daraei A, Pieters M, Baker SR, et al. Automated fiber diameter and porosity measurements of plasma clots in scanning electron microscopy images. Biomolecules. 2021;11(10):1536.

9. Kouwer PH, Koepf M, Le Sage VA, et al. Responsive biomimetic networks from polyisocyanopeptide hydrogels. Nature. 2013;493(7434):651–655.

10. Hell SW, Wichmann J. Breaking the diffraction resolution limit by stimulated emission: stimulated-emission-depletion fluorescence microscopy. Opt Lett. 1994;19(11):780–782.

11. Stojanov S, Berlec A. Electrospun nanofibers as carriers of microorganisms, stem cells, proteins, and nucleic acids in therapeutic and other applications. Front Bioeng Biotechnol.2020;8:130.

12. Ho SY, Chao CY, Huang HL, Chiu TW, Charoenkwan P, Hwang E. NeurphologyJ: An automatic neuronal morphology quantification method and its application in pharmacological discovery. BMC Bioinform. 2011;12.

13. Bridge KI, Philippou H, Ariens R. Clot properties and cardiovascular disease. Thromb Haemost.2014;112(5):901–908.

14. Dauwerse S, Ten Cate H, Spronk HMH, Nagy M. The composition and physical properties of clots in COVID-19 pathology. Diagnostics (Basel). 2022;12(3):580.

15. de Vries JJ, Visser C, Geers L, et al. Altered fibrin network structure and fibrinolysis in intensive care unit patients with COVID-19, not entirely explaining the increased risk of thrombosis. J Thromb Haemost. 2022;20(6):1412–1420.

16. Weisel JW, Litvinov RI. Fibrin formation, structure and properties. Subcell Biochem.2017;82:405–456.

17. Undas A. Fibrin clot properties and their modulation in thrombotic disorders. Thromb Haemost. 2014;112(1):32–42.

18. Undas A, Ariens RA. Fibrin clot structure and function: a role in the pathophysiology of arterial and venous thromboembolic diseases. Arterioscler Thromb Vasc Biol. 2011;31(12):e88–99.

19. Collet JP, Park D, Lesty C, et al. Influence of fibrin network conformation and fibrin fiber diameter on fibrinolysis speed: dynamic and structural approaches by confocal microscopy. Arterioscler Thromb Vasc Biol. 2000;20(5):1354–1361.

20. Stevens CR, Berenson J, Sledziona M, Moore TP, Dong L, Cheetham J. Approach for semi-automated measurement of fiber diameter in murine and canine skeletal muscle. PLoS ONE.2020;15(12):e0243163.

21. Zong Y, Pruner I, Antovic A, et al. Phosphatidylserine positive microparticles improve hemostasis in in-vitro hemophilia A plasma models. Sci Rep. 2020;10(1):7871.

22. Canver AC, Morss Clyne A. Quantification of multicellular organization, junction integrity, and substrate features in collective cell migration. Microsc Microanal. 2017;23(1):22–33.

23. Stein AM, Vader DA, Jawerth LM, Weitz DA, Sander LM. An algorithm for extracting the network geometry of three-dimensional collagen gels. J Microsc. 2008;232(3):463–475.

24. Bredfeldt JS, Liu YM, Pehlke CA, et al. Computational segmentation of collagen fibers from second-harmonic generation images of breast cancer. J Biomed Opt. 2014;19(1).

25. Jaeschke A, Eckert H, Bray LJ. Qiber3D-An open-source software package for the quantitative analysis of networks from 3D image stacks. GigaScience. 2022;11:giab091.

26. Liu Z, Speroni L, Quinn KP, et al. 3D organizational mapping of collagen fibers elucidates matrix remodeling in a hormone-sensitive 3D breast tissue model. Biomaterials. 2018;179:96–108.

27. Rezakhaniha R, Agianniotis A, Schrauwen JT, et al. Experimental investigation of collagen waviness and orientation in the arterial adventitia using confocal laser scanning microscopy. Biomech Model Mechanobiol. 2012;11(3-4):461–473.

28. Liu Y, Keikhosravi A, Mehta GS, Drifka CR, Eliceiri KW. Methods for quantifying fibrillar collagen alignment. Methods Mol Biol. Vol. 1627; 2017:429–451.

29. Persson NE, McBride MA, Grover MA, Reichmanis E. Automated Analysis of Orientational Order in Images of Fibrillar Materials. Chem Mater. 2017;29(1):3–14.

30. Kartasalo K, Pölönen R-P, Ojala M, et al. CytoSpectre: a tool for spectral analysis of oriented structures on cellular and subcellular levels. BMC Bioinform. 2015;16(1):344.

31. van Haaften EE, Wissing TB, Rutten MCM, et al. Decoupling the Effect of Shear Stress and Stretch on Tissue Growth and Remodeling in a Vascular Graft. Tissue Eng Part C Methods.2018;24(7):418–429.

32. Morrill EE, Tulepbergenov AN, Stender CJ, Lamichhane R, Brown RJ, Lujan TJ. A validated software application to measure fiber organization in soft tissue. Biomech Model Mechanobiol.2016;15(6):1467–1478.

33. Marcotti S, de Freitas DB, Troughton LD, et al. A workflow for rapid unbiased quantification of fibrillar feature alignment in biological images. Front Comput Sci. 2021;3.

34. Vos BE, Martinez-Torres C, Burla F, Weisel JW, Koenderink GH. Revealing the molecular origins of fibrin’s elastomeric properties by in situ X-ray scattering. Acta Biomater. 2020;104:39–52.

35. Hotaling NA, Bharti K, Kriel H, Simon CG. DiameterJ: A validated open source nanofiber diameter measurement tool. Biomaterials. 2015;61:327–338.

36. Hildebrand T, Ruegsegger P. Quantification of Bone Microarchitecture with the Structure Model Index. Comput Methods Biomech Biomed Engin. 1997;1(1):15–23.

37. Dougherty D, Kunzelmann K. Computing local thickness of 3D structure with ImageJ. Micros Microanal. 2007;13:1678–1679.

38. Corliss BA, Doty RW, Mathews C, Yates PA, Zhang T, Peirce SM. REAVER: A program for improved analysis of high-resolution vascular network images. Microcirculation. 2020;27(5):e12618.

39. Fricker M, Heaton L, Jones N, Obara B, Müller SJ, Meyer AJ. Quantitation of ER structure and function. Methods Mol Biol. Vol. 1691; 2018:43–66.

40. Dabbah MA, Graham J, Petropoulos I, Tavakoli M, Malik RA. Dual-model automatic detection of nerve-fibres in corneal confocal microscopy images. International Conference on Medical Image Computing and Computer-Assisted Intervention. 2010;13(Pt 1):300–307.

41. Chen X, Graham J, Dabbah MA, Petropoulos IN, Tavakoli M, Malik RA. An automatic tool for quantification of nerve fibers in corneal confocal microscopy images. IEEE Trans Biomed Eng.2017;64(4):786–794.

42. Shkarin R, Shkarin A, Shkarina S, et al. Quanfima: An open source Python package for automated fiber analysis of biomaterials. PLoS ONE. 2019;14(4):e0215137.

43. Murphy R, Turcott A, Banuelos L, Dowey E, Goodwin B, Cardinal KO. SIMPoly: a matlab-based image analysis tool to measure electrospun polymer scaffold fiber diameter. Tissue Eng Part C Methods. 2020;26(12):628–636.

44. Hotaling NA, Bharti K, Kriel H, Simon CG, Jr. Dataset for the validation and use of DiameterJ an open source nanofiber diameter measurement tool. Data Brief. 2015;5:13–22.

45. Shah GA, Ferguson IA, Dhall TZ, Dhall DP. Polydispersion in the diameter of fibers in fibrin networks: consequences on the measurement of mass-length ratio by permeability and turbidity. Biopolymers. 1982;21(6):1037–1047.

46. Zudaire E, Gambardella L, Kurcz C, Vermeren S. A computational tool for quantitative analysis of vascular networks. PLoS ONE. 2011;6(11):e27385.

47. Xu T, Vavylonis D, Tsai FC, et al. SOAX: a software for quantification of 3D biopolymer networks. Sci Rep. 2015;5:9081.

48. Vecchio DA, Mahler SH, Hammig MD, Kotov NA. Structural analysis of nanoscale network materials using graph theory. ACS Nano. 2021;15(8):12847–12859.

49. Cross SS. Fractals in pathology. J Pathol. 1997;182(1):1–8.

50. Doube M, Kłosowski MM, Arganda-Carreras I, et al. BoneJ: Free and extensible bone image analysis in ImageJ. Bone. 2010;47(6):1076–1079.

51. Hood JE, Yesudasan S, Averett RD. Glucose concentration affects fibrin clot structure and morphology as evidenced by fluorescence imaging and molecular simulations. Clin Appl Thromb Hemost. 2018;24(9_suppl):104S–116S.

52. Münster S, Fabry B. A simplified implementation of the bubble analysis of biopolymer network pores. Biophys J. 2013;104(12):2774–2775.

53. Krauss P, Metzner C, Lange J, Lang N, Fabry B. Parameter-free binarization and skeletonization of fiber networks from confocal image stacks. PLoS ONE. 2012;7(5):e36575.

54. Jamshidi M, Falamaki C. Image analysis method for heterogeneity and porosity characterization of biomimetic hydrogels. F1000Res. 2020;9:1461.

55. Wang S, Zhong S, Lim CT, Nie H. Effects of fiber alignment on stem cells-fibrous scaffold interactions. J Mater Chem B. 2015;3(16):3358–3366.

56. Laurens N, Koolwijk P, de Maat MP. Fibrin structure and wound healing. J Thromb Haemost.2006;4(5):932–939.

57. Campbell RA, Aleman M, Gray LD, Falvo MR, Wolberg AS. Flow profoundly influences fibrin network structure: implications for fibrin formation and clot stability in haemostasis. Thromb Haemost. 2010;104(6):1281–1284.

58. Tutwiler V, Maksudov F, Litvinov RI, Weisel JW, Barsegov V. Strength and deformability of fibrin clots: Biomechanics, thermodynamics, and mechanisms of rupture. Acta Biomater.2021;131:355–369.

59. Alvarado J, Mulder BM, Koenderink GH. Alignment of nematic and bundled semiflexible polymers in cell-sized confinement. Soft Matter. 2014;10(14):2354–2364.

60. Narayanan G, Tekbudak MY, Caydamli Y, Dong J, Krause WE. Accuracy of electrospun fiber diameters: The importance of sampling and person-to-person variation. Polym Test. 2017;61:240–248.

61. Ryan EA, Mockros LF, Weisel JW, Lorand L. Structural origins of fibrin clot rheology. Biophys J.1999;77(5):2813–2826.

62. Hawkins K, Badiei N, Weisel J, et al. Fractal dimension: a biomarker for detecting acute thromboembolic disease. Critical Care. 2012;16(1):P431.

63. Lawrence MJ, Sabra A, Thomas P, et al. Fractal dimension: a novel clot microstructure biomarker use in ST elevation myocardial infarction patients. Atherosclerosis. 2015;240(2):402–407.

64. Davies NA, Harrison NK, Morris RH, et al. Fractal dimension (df) as a new structural biomarker of clot microstructure in different stages of lung cancer. Thromb Haemost. 2015;114(6):1251–1259.

65. Bossler F, Maurath J, Dyhr K, Willenbacher N, Koos E. Fractal approaches to characterize the structure of capillary suspensions using rheology and confocal microscopy. J Rheol. 2018;62(1):183–196.

66. Fischer T, Hayn A, Mierke CT. Fast and reliable advanced two-step pore-size analysis of biomimetic 3D extracellular matrix scaffolds. Sci Rep. 2019;9(1):8352.

67. Nakamura K, Suda T, Matsumoto K. Characterization of pore size distribution of non-woven fibrous filter by inscribed sphere within 3D filter model. Sep Purif Technol. 2018;197:289–294.

68. In Hwan S, Young Jun C, Chang Kyu P. Automatic volumetric measurement of nanofiber webs using metaball approximation based on scanning electron microscope images. Text Res J.2009;80(11):995–1003.

69. Lang NR, Munster S, Metzner C, et al. Estimating the 3D pore size distribution of biopolymer networks from directionally biased data. Biophys J. 2013;105(9):1967–1975.

70. Moher D, Liberati A, Tetzlaff J, Altman DG, Group P. Preferred reporting items for systematic reviews and meta-analyses: the PRISMA statement. J Clin Epidemiol. 2009;62(10):1006–1012.

71. de Maat MP, van Schie M, Kluft C, Leebeek FW, Meijer P. Biological variation of hemostasis variables in thrombosis and bleeding: consequences for performance specifications. Clin Chem.2016;62(12):1639–1646.

