## Supplementary for "A systematic review and comparison of automated tools for quantification of fibrous networks"

- Supplementary Figures I-VIII
- Supplementary Tables I-XVIII
- Supplementary References
- Literature search terms
- Python code for simulated fibrous networks
- ZIP file of used confocal, STED, SEM, and simulated images
- Dataset results tools

### Supplementary Figures

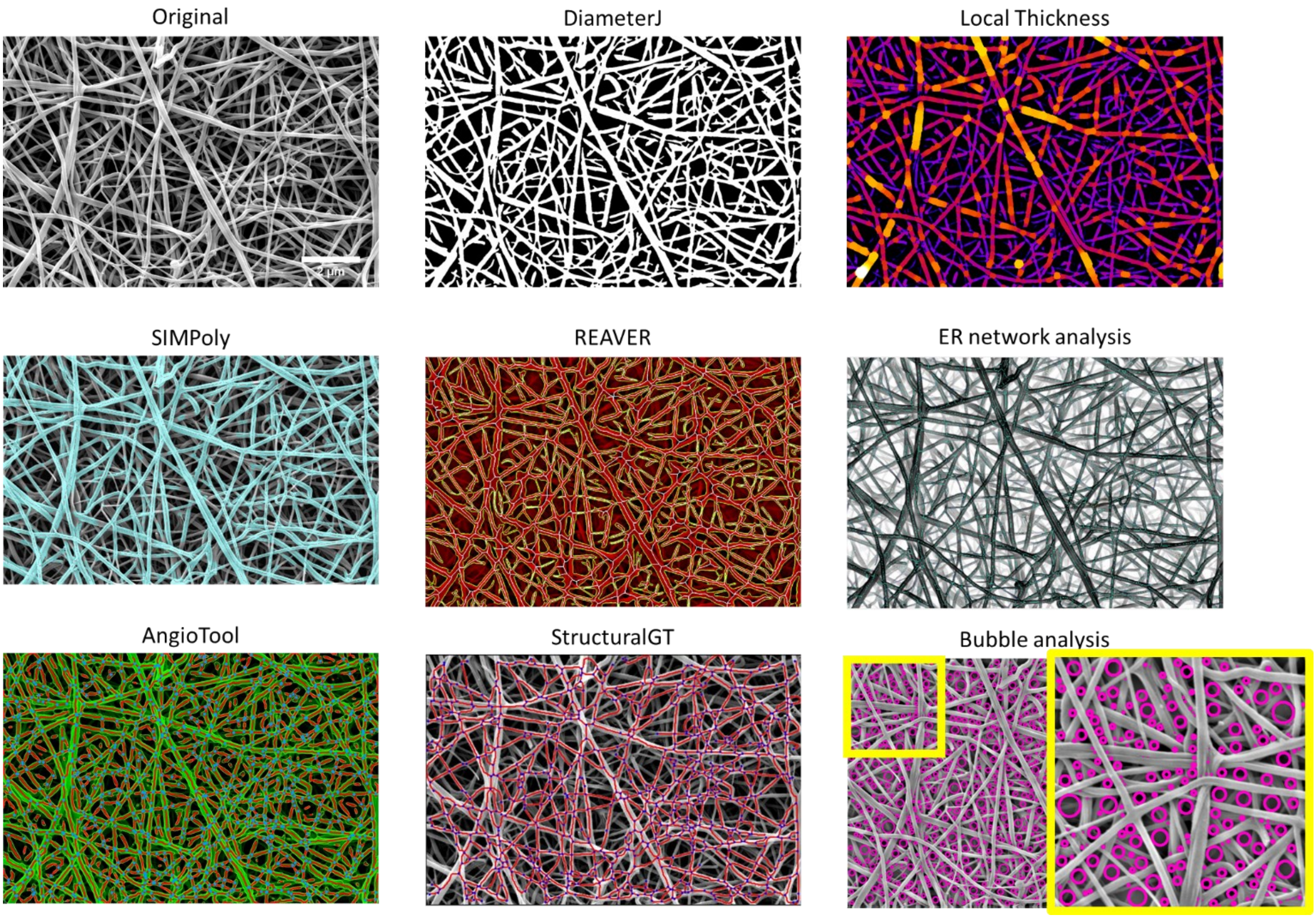


**Supplementary Figure I**. Examples of the automated tools applied to the same SEM image. The first image shows the original SEM image, with subsequently the visual representations of the results of the different tools. For DiameterJ, the image after segmentation is shown with fibers in white and background in black. For the Local Thickness plugin, fibers are color-coded with fibers in yellow indicating higher thickness than fibers depicted in purple. SIMPoly indicates the identified fibers in light blue. REAVER indicates the center of the fibers with a white line, while the borders are indicated using yellow lines. The ER network analysis indicates center of fibers using light blue lines, and fibers are shown in black on a white background. AngioTool shows fibers in green, with a red line indicating the center of the fibers and blue dots indicating branch points. StructuralGT presents the centers of the fibers by red lines with blue dots indicating branch points. Finally, pores identified by the Bubble analysis were depicted in pink on the original image. For the Bubble analysis, we show a zoomed-in view of the yellow box for clarity. No visual representation was available of the application of BoneJ, Algorithms Hood (fractal dimension) and Quanfima (porosity), since these tools do not provide this output.

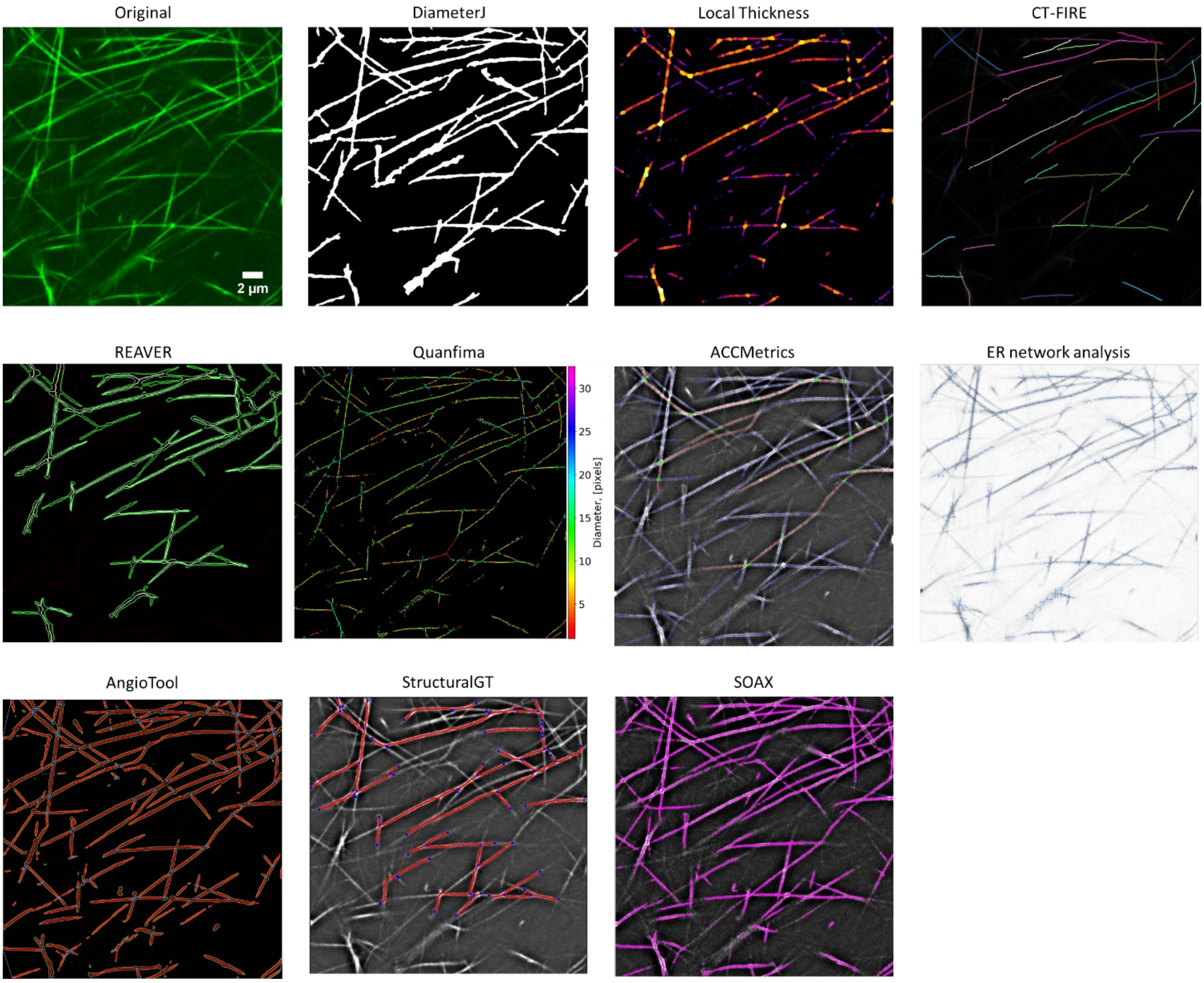


**Supplementary Figure II**. Examples of the automated tools applied to the same STED image. The first image shows the original STED image, with subsequently the visual representations of the results of the different tools. For DiameterJ, the image after segmentation is shown with fibers in white and background in black. For the Local Thickness plugin, fibers are color-coded with fibers in yellow indicating higher thickness than fibers depicted in purple. CT-FIRE shows the identified fibers in different colors depicting separate fibers. REAVER indicates the center of the fibers with a white line, while the borders are indicated using yellow lines. Quanfima shows the identified fibers with a color-coding based on the diameter, with red/yellow meaning smaller diameters and green/blue meaning higher diameters. ACCMetrics depicts the identified fibers using blue and red lines with green dots representing branch points. The ER network analysis indicates center of fibers using light blue lines, and fibers are shown in grey on a white background. AngioTool shows fibers in green, with a red line indicating the center of the fibers and blue dots indicating branch points. StructuralGT presents the centers of the identified fibers by red lines with blue dots indicating branch points. SOAX shows fibers in purple on a black background. No visual representation was available of the application of BoneJ and Algorithms Hood (fractal dimension), since these tools do not provide this output.

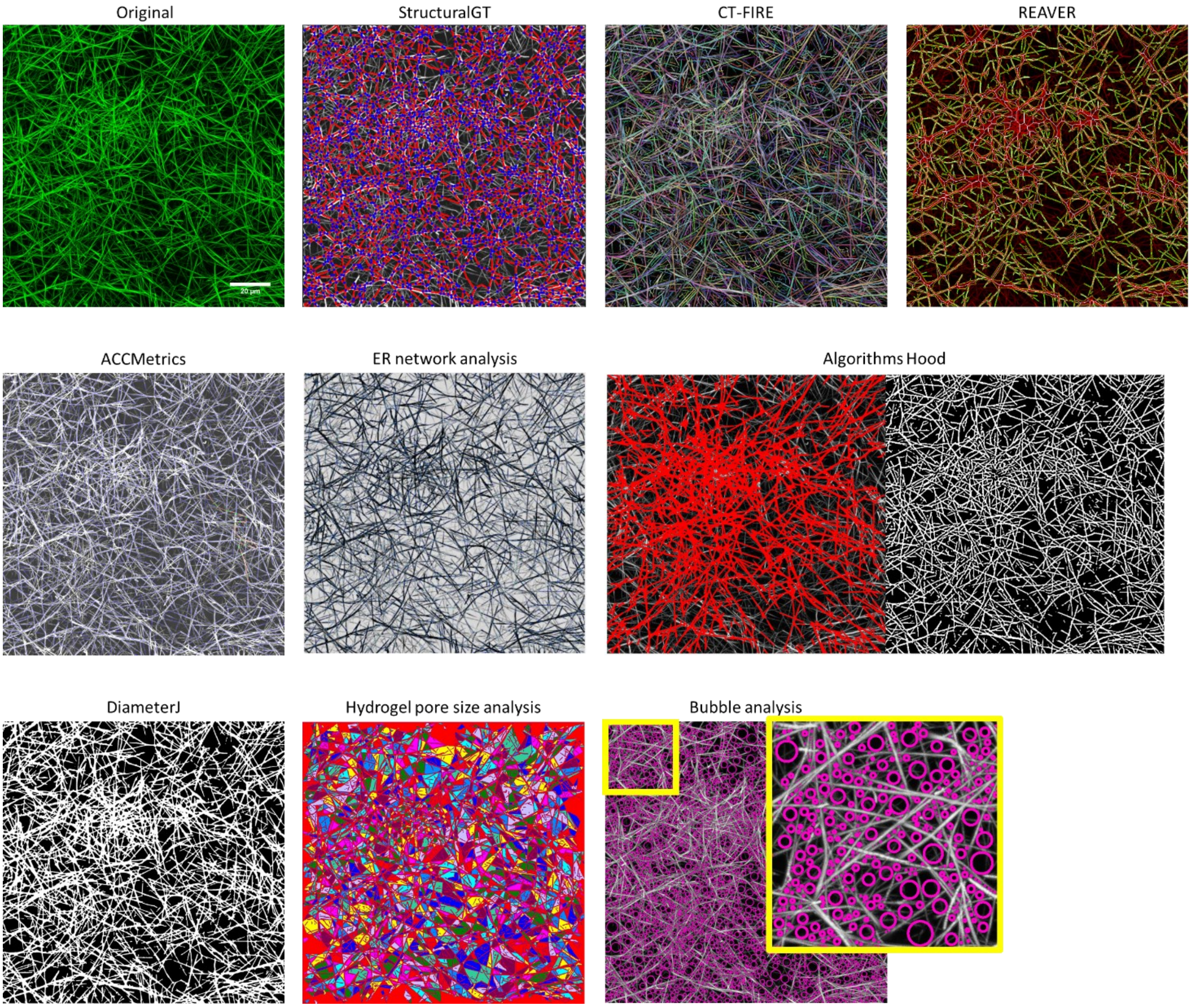


**Supplementary Figure III**. Examples of the automated tools applied to the same confocal image of a fibrin network formed under static conditions. The first image shows the original confocal image, with subsequently the visual representations of the results of the different tools. StructuralGT presents the centers of the fibers by red lines with blue dots indicating branch points. CT-FIRE shows the identified fibers in different colors depicting separate fibers. REAVER indicates the center of the fibers with a white line, while the borders are indicated using yellow lines. ACCMetrics depicts the identified fibers using blue and red lines with green dots representing branch points. The ER network analysis indicates center of fibers using blue lines, and fibers are shown in black on a white background. Algorithms by Hood et al. show identified fibers in red over the original image and identified fibers in white on a black background. For DiameterJ, the image after segmentation is shown with fibers in white and background in black. The Hydrogel pore size analysis depicts separate pores in different colors with fibers and background in red. Finally, pores identified by the Bubble analysis were depicted in pink on the original image. For the Bubble analysis, we show a zoomed-in view of the yellow box for clarity. No visual representation was available of the application of BoneJ, Pore size analysis (Krauss), and Quanfima (porosity), since these tools do not provide this output.

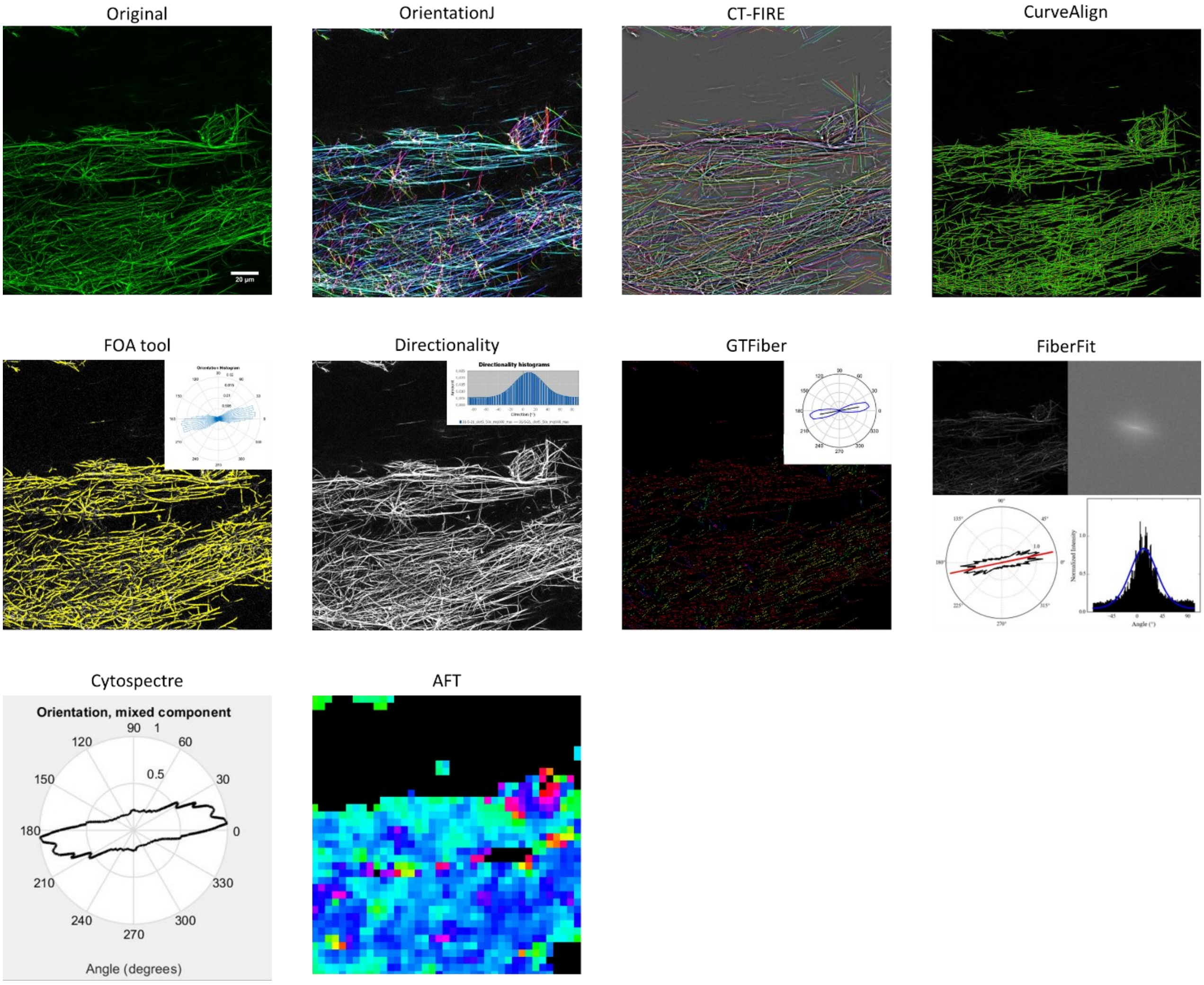


**Supplementary Figure IV**. Examples of the automated tools applied to the same confocal image of a fibrin network formed under flow using a shear rate of 50 $s^{-1}$. The first image shows the original confocal image with subsequently the visual representations of the results of the different tools. OrientationJ depicts identified fibers using color-coding regarding the orientation of the fibers. CT-FIRE shows the identified fibers in different colors depicting separate fibers. CurveAlign depicts identified fibers in green. The FOA tool shows identified fibers in yellow, with the corresponding orientation histogram of the fibers in the right upper corner of the image. The Directionality plugin depicts fibers in white on a black background with the corresponding orientation histogram in the right upper corner of the image. GTFiber shows identified fibers in colors representing different orientations with the orientation distribution in the right upper corner of the image. The black line indicates the average orientation and the blue line indicates the count of pixels of a given orientation. FiberFit shows the original image in the left upper corner of the panel, the fast Fourier Transform of the image in the right upper corner, the fiber orientation distribution in black with the average orientation in red in the left lower corner, and the orientation histogram in the right lower corner. Cytospectre does not result in a visual representation of the fibers, but does show the orientation distribution as a black line per degree of orientation. AFT returns an heatmap depicting the orientations of local alignment vectors in the image with different colors depicting different orientations.

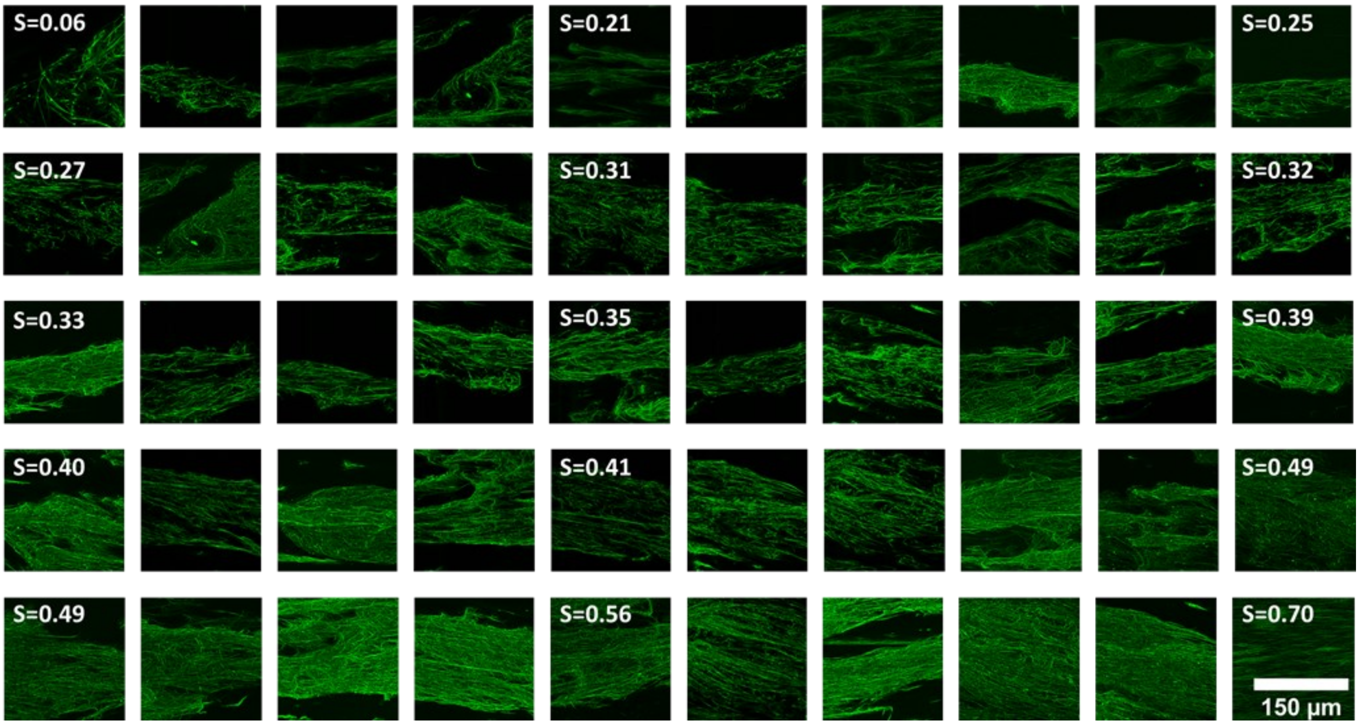


**Supplementary Figure V.** Overview of the 50 confocal images of fibrin fibers formed under flow ordered from low to high degree of fiber alignment based on their corresponding nematic order parameters (S) as calculated by OrientationJ.

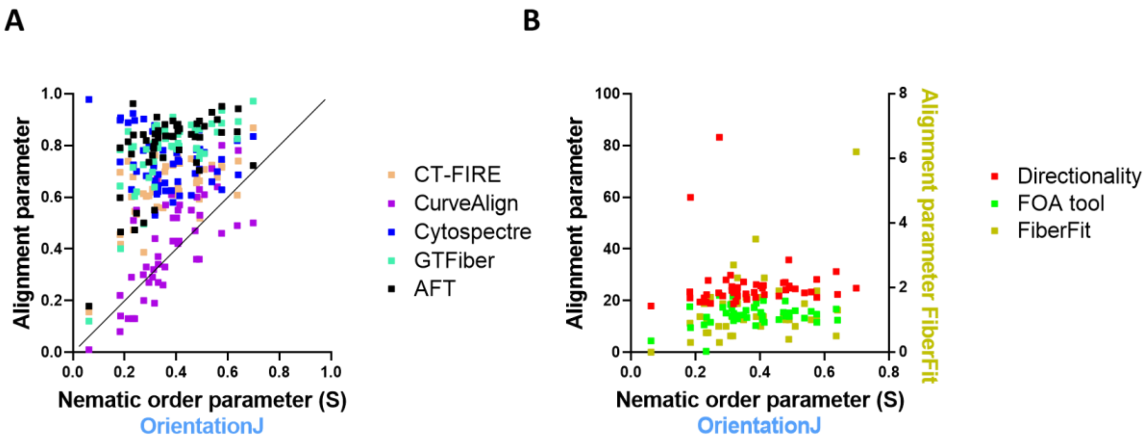


**Supplementary Figure VI.** Correlation between the results of the different tools used to quantify fibrin fiber alignment. Shear rates ranged between 50 $s^{-1}$ and 250 $s^{-1}$. Every dot represents one image. The nematic order parameter (S) quantified by OrientationJ was placed on the X-axis, while the results of the other tools are placed on the Y-axis. OrientationJ was used as reference because of its proven sensitivity for fiber alignment[1]. A) Correlation between OrientationJ and the other tools resulting in an order parameter between 0 and 1. The solid line indicates the x=y-line. B) Correlation between OrientationJ and the tools resulting in a parameter describing the spread of the orientation distribution.

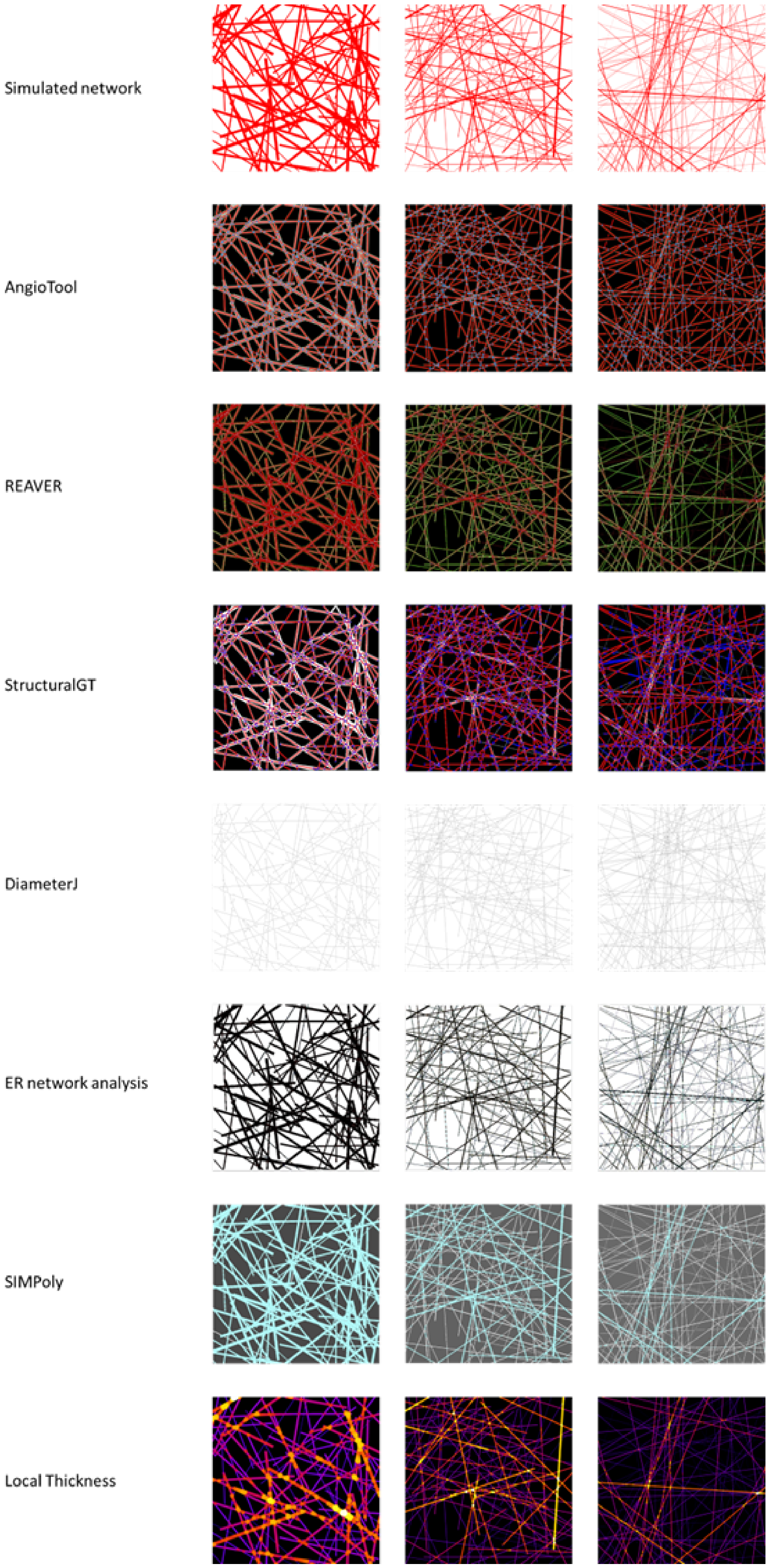


**Supplementary Figure VII**. Visual presentations of the automated tools applied to three simulated fibrous networks with 100 fibers with known mean diameter (22, 7.4, and 3.6 pixels, respectively), total fiber length (94620, 141720, and 178600 pixels, respectively), and number of branch points (745, 1654, and 2531, respectively). The images on the first row depict the simulated networks with the fibers in red on a white background. The subsequent rows show the results of the different tools for the three simulated networks. AngioTool shows fibers in green, with a red line indicating the center of the fibers and blue dots indicating branch points. REAVER indicates the center of the fibers with a white line, while the borders are indicated using yellow lines. StructuralGT presents the centers of the fibers by red lines with blue dots indicating branch points. For DiameterJ, centerlines of identified fibers are depicted in black on a white background. The ER network analysis shows identified fibers in black on a white background. SIMPoly indicates the identified fibers in light blue. For the Local Thickness plugin, fibers are color-coded with fibers in yellow indicating higher thickness than fibers depicted in purple.

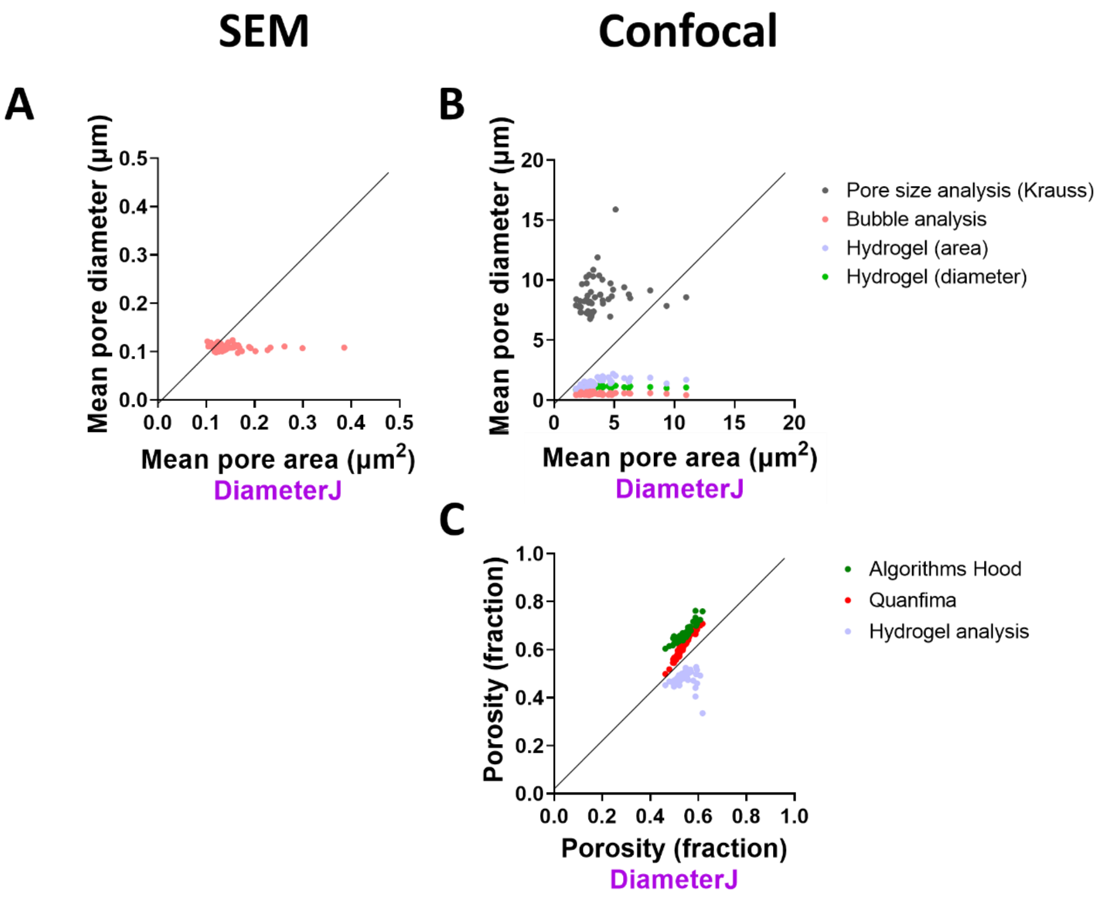


**Supplementary Figure VIII.** Correlation between the results of the different tools used to quantify the pore size (A) or porosity (B) in SEM and confocal images. The solid lines indicate the x=y-lines.

### Supplementary Tables

**Supplementary Table I.** An overview of excluded image analysis tools with reasons

| No quantification of (relevant) parameters (n=12) | Requires specific instrument or paid software (n=13) | Manual steps involved (n=5) | Other imaging (n=5) | Not suitable for our images of fibrin (n=13) |
|---|---|---|---|---|
| ANNA-PALM[2] | Amira software Xtracing[3] | FiberApp[4] | FiberScout[5] | ARGSLab[6] |
| Ilastik[7] | PAQXOS[8] | FibrilTool[9] | Angiogenesis Analyzer[10] | FibrilJ[11] |
| Ilastik + MATLAB script[12] | Fiber Thickness App[13] | NeuronJ[14] | Cytoseg[15] | MTrack[16] |
| JFilament[17] | Phenom FiberMetric Software[18] | Continuity software[19] | PyFibre[20] | MyoSAT[21] |
| Adaptive Resolution Orientation Space[22] | Morfi Analyzer[23] | Monte Carlo simulation[24] | QuantAn[25] | Neurphology[26] |
| Orientation aware neural network[27] | SEMAnalyzer[28] | | | ThunderSTORM[29] |
| Fiber Finding Algorithm[30] | FibreQuant[28] | | | Vessel Analysis Plugin[31] |
| FiberQ[32] | CellArchitect[33] | | | Skeleton analysis plugin[34] |
| Image-Based Fiber Orientation and Alignment Calculator[35] | FibroIndex[36] | | | ecm MATLAB script[37]<br>TSOAX[38] |
| SFEX[39] | Imaris | | | Fiber Analysis Algorithm[40] |
| SIFNE[41] | Filament network tracing algorithm FiNTA[42] | | | KerNet[43] |
| Persistence length analyser[44] | Image Pro[45]<br>3D profilometer[47] | | | TWOMBLI[46] |

**Supplementary Table II**. Mean and median of alignment parameters and the correlations between different tools

| Name tool | Mean ± SD | Median [25th-75th percentile] | OrientationJ | CurveAlign | GTFiber | CT-FIRE | AFT | Cytospectre | FiberFit | Directionality | FOA tool |
|---|---|---|---|---|---|---|---|---|---|---|---|
| **OrientationJ** | 0.37 ± 0.13 | 0.35 [0.28-0.48] | | 0.73* | 0.65* | 0.60* | 0.53* | -0.27 | 0.36* | -0.06 | 0.25 |
| **CurveAlign** | 0.41 ± 0.19 | 0.43 [0.29-0.55] | 0.73* | | 0.63* | 0.46* | 0.62* | -0.65* | 0.32* | -0.25 | 0.09 |
| **GTFiber** | 0.77 ± 0.14 | 0.80 [0.75-0.85] | 0.65* | 0.63* | | 0.87* | 0.88* | -0.63* | 0.55* | -0.40* | 0.17 |
| **CT-FIRE** | 0.63 ± 0.11 | 0.63 [0.60-0.72] | 0.60* | 0.46* | 0.87* | | 0.74* | -0.46* | 0.57* | -0.37* | 0.09 |
| **AFT** | 0.79 ± 0.15 | 0.83 [0.76-0.87] | 0.53* | 0.62* | 0.88* | 0.74* | | -0.59* | 0.28 | -0.39* | -0.01 |
| **Cytospectre** | 0.74 ± 0.10 | 0.73 [0.66-0.80] | -0.27 | -0.65* | -0.63* | -0.46* | -0.59* | | -0.47* | 0.40* | -0.01 |
| **FiberFit** | 1.32 ± 0.95 | 1.10 [0.80-1.63] | 0.36* | 0.32* | 0.55* | 0.57* | 0.28 | -0.47* | | -0.29* | -0.16 |
| **Directionality** | 25.8 ± 10.3 | 24.8 [19.5-35.8] | -0.06 | -0.25 | -0.40* | -0.37* | -0.39* | 0.40* | -0.29* | | 0.31* |
| **FOA tool** | 14.3 ± 3.9 | 14.3 [12.7-16.5] | 0.25 | 0.09 | 0.17 | 0.09 | -0.01 | -0.01 | -0.16 | 0.31* | |

Pearson correlation coefficient; *$p<0.05$. Green shading indicates significant association and no shading (white) indicates no significant association. OrientationJ gives the nematic order parameter (S), CurveAlign the alignment coefficient, GTfiber gives the orientational order parameter ($S_{full}$), CT-FIRE gives a parameter termed Alignment, AFT results in a median order parameter, Cytospectre reports the circular variance, FiberFit returns the fiber dispersion parameter, Directionality the dispersion, and the FOA tool gives the standard deviation of the histogram.

**Supplementary Table III**. Quantification of fiber alignment in synthetic images with known dispersion parameters

| Known dispersion | OrientationJ | CurveAlign | GTFiber | CT-FIRE | AFT | Cytospectre | FiberFit | FOA tool |
|---|---|---|---|---|---|---|---|---|
| 0.20 | 0.07 | 0.01 | 0.01 | 0.02 | 0.38 | 0.88 | 0.27 | NA |
| 0.30 | 0.02 | 0.02 | 0.07 | 0.00 | 0.34 | 0.86 | 0.38 | 54.92 |
| 0.40 | 0.01 | 0.09 | 0.09 | 0.01 | 0.44 | 0.83 | 0.46 | 52.69 |
| 0.50 | 0.02 | 0.11 | 0.17 | 0.01 | 0.38 | 0.79 | 0.53 | 51.10 |
| 0.60 | 0.06 | 0.14 | 0.25 | 0.07 | 0.34 | 0.72 | 0.66 | 49.71 |
| 0.70 | 0.10 | 0.12 | 0.32 | 0.18 | 0.31 | 0.77 | 0.73 | 47.95 |
| 0.80 | 0.07 | 0.18 | 0.23 | 0.28 | 0.40 | 0.65 | 0.81 | 44.37 |
| 0.90 | 0.11 | 0.18 | 0.34 | 0.28 | 0.42 | 0.60 | 0.96 | 42.33 |
| 1.00 | 0.15 | 0.24 | 0.46 | 0.37 | 0.43 | 0.62 | 1.06 | 39.26 |
| 2.00 | 0.32 | 0.62 | 0.67 | 0.57 | 0.65 | 0.35 | 2.17 | 27.46 |
| 3.00 | 0.44 | 0.81 | 0.77 | 0.72 | 0.79 | 0.26 | 2.95 | 21.46 |
| 4.00 | 0.39 | 0.86 | 0.78 | 0.80 | 0.87 | 0.23 | 4.08 | 19.05 |
| 5.00 | 0.45 | 0.89 | 0.83 | 0.84 | 0.90 | 0.19 | 5.07 | 16.61 |
| 10.00 | 0.60 | 0.96 | 0.92 | 0.91 | 0.96 | 0.11 | 10.17 | 11.13 |
| **Correlation with known dispersion** | 0.85* | 0.86* | 0.83* | 0.85* | 0.87* | -0.87* | 0.99* | -0.73* |

The first column gives the known fiber dispersion parameter and the subsequent columns report the results from the automated analysis tools. Values are color-coded within their column: darker red means more aligned networks, darker green means less aligned networks. The bottom row reports Pearsons correlation coefficients for the correlation between the known dispersion values and the results of the automated analysis tools. *p<0.05. NA, not available.

**Supplementary Table IV**. Mean and median of fiber diameter and the correlations between different tools in SEM images

| Name tool | Mean diameter ± SD (nm) | Median [25th-75th percentile] (nm) | Manual | Local Thickness | DiameterJ | SIMPoly | REAVER | ER network analysis |
|---|---|---|---|---|---|---|---|---|
| **Manual** | 134.9 ± 22.9 | 128.8 [120.4-143.9] | | -0.16 | 0.90* | 0.92* | 0.39* | 0.82* |
| **Local Thickness** | 140.9 ± 50.0 | 135.7 [101.3-181.6] | -0.16 | | -0.21 | -0.36* | 0.44* | -0.06 |
| **DiameterJ** | 111.5 ± 10.7 | 108.5 [104.0-114.8] | 0.90* | -0.21 | | 0.92* | 0.27 | 0.78* |
| **SIMPoly** | 147.0 ± 21.3 | 141.1 [132.7-153.2] | 0.92* | -0.36* | 0.92* | | 0.21 | 0.77* |
| **REAVER** | 176.3 ± 28.3 | 169.7 [156.4-192.5] | 0.39* | 0.44* | 0.27 | 0.21 | | 0.42* |
| **ER network analysis** | 134.4 ± 20.8 | 125.8 [119.2-144.1] | 0.82* | -0.06 | 0.78* | 0.77* | 0.42* | |

Pearson correlation coefficient; *p<0.05. Green shading indicates significant association, orange shading indicates significant association in the opposite direction of what we expected, and no shading (white) indicates no significant association.

**Supplementary Table V**. Mean and median of fiber diameter and the correlations between different tools in STED images

| Name tool | Mean diameter ± SD (nm) | Median [25th-75th percentile] (nm) | ER network analysis | ACCMetrics | Local Thickness | DiameterJ | CT-FIRE | REAVER | Quanfima |
|---|---|---|---|---|---|---|---|---|---|
| **ER network analysis** | 279.4 ± 67.9 | 275.6 [248.2-309.4] | | 0.70* | 0.89* | 0.55* | 0.62* | 0.83* | 0.51* |
| **ACCMetrics** | 229.9 ± 11.6 | 228.0 [221.2-235.7] | 0.70* | | 0.88* | 0.93* | 0.74* | 0.77* | 0.82* |
| **Local Thickness** | 278.1 ± 51.8 | 266.1 [244.9-303.5] | 0.89* | 0.88* | | 0.86* | 0.69* | 0.84* | 0.75* |
| **DiameterJ** | 311.5 ± 42.3 | 310.7 [273.1-325.4] | 0.55* | 0.93* | 0.86* | | 0.69* | 0.69* | 0.82* |
| **CT-FIRE** | 201.1 ± 22.8 | 197.0 [183.6-215.6] | 0.62* | 0.74* | 0.69* | 0.69* | | 0.73* | 0.51* |
| **REAVER** | 416.8 ± 109.8 | 402.0 [342.2-477.1] | 0.83* | 0.77* | 0.84* | 0.69* | 0.73* | | 0.78* |
| **Quanfima** | 254.5 ± 31.9 | 250.6 [232.4-264.0] | 0.51* | 0.82* | 0.75* | 0.82* | 0.51* | 0.78* | |

Pearson correlation coefficient; *p<0.05. Green shading indicates significant association and no shading (white) indicates no significant association.

**Supplementary Table VI**. Quantification of fiber diameters in synthetic images and simulated networks with known (mean) diameters in pixels

| **Constant diameter (pixels)** | **DiameterJ** | **Local Thickness** | **SIMPoly** | **Quanfima** | **ER network analysis** | **REAVER** |
|---|---|---|---|---|---|---|
| 5 | 6.06 | 9.96 | 12.98 | 9.19 | 7.20 | 30.55 |
| 5 | 5.94 | 8.30 | 12.30 | 11.49 | 6.85 | 15.33 |
| 5 | 5.89 | 8.09 | 12.01 | 11.99 | 6.75 | 14.10 |
| 10 | 10.22 | 15.90 | 9.94 | 14.38 | 11.98 | 36.25 |
| 10 | 10.17 | 13.61 | 9.56 | 17.66 | 10.92 | 21.88 |
| 10 | 10.15 | 13.82 | 9.56 | 18.05 | 10.92 | 20.91 |
| 15 | 15.63 | 23.89 | 15.57 | 16.55 | 18.29 | 31.63 |
| 15 | 15.65 | 23.88 | 15.61 | 17.31 | 18.21 | NA |
| 15 | 15.66 | 23.33 | 15.41 | 17.44 | 17.97 | 29.18 |
| 25 | 25.39 | 44.02 | 25.03 | 17.79 | 29.45 | 44.08 |
| 25 | 25.45 | 35.00 | 25.00 | 20.43 | 26.91 | 35.90 |
| 25 | 25.44 | 36.05 | 25.06 | 20.54 | 27.08 | 35.92 |
| 50 | 49.72 | 83.77 | 49.75 | 20.90 | 36.46 | 76.01 |
| 50 | 49.79 | 71.09 | 49.60 | 23.23 | 35.31 | 64.03 |
| 50 | 49.68 | 69.67 | 49.63 | 22.51 | 34.64 | 62.54 |

| **Average diameter (pixels)** | **DiameterJ** | **Local Thickness** | **SIMPoly** | **Quanfima** | **ER network analysis** | **REAVER** |
|---|---|---|---|---|---|---|
| 20 | 9.94 | 63.19 | | 18.48 | 17.27 | 57.06 |
| 20 | 17.14 | 36.37 | 23.65 | 17.44 | 24.75 | 39.77 |
| 20 | 16.32 | 51.08 | 16.26 | 15.72 | 23.03 | 56.08 |
| 35 | 28.76 | 61.18 | 28.50 | 19.59 | 29.89 | 57.59 |
| 35 | 22.67 | 86.79 | 21.99 | 19.45 | 27.23 | 69.29 |
| 35 | 17.98 | 67.57 | 45.11 | 19.87 | 29.42 | 58.53 |
| 50 | 55.76 | 88.73 | 55.67 | 21.18 | 33.35 | 78.80 |
| 50 | 35.90 | 102.50 | 63.13 | 20.20 | 31.18 | 87.14 |

| | | | | | | |
|---|---|---|---|---|---|---|
| 50 | 32.62 | 106.81 | 55.33 | 19.50 | 32.40 | 89.17 |
| **Simulated networks: mean diameter (pixels)** | **DiameterJ** | **Local Thickness** | **SIMPoly** | **Quanfima** | **ER network analysis** | **REAVER** |
| 22 | 20.9 | 33.4 | 23.1 | NA | 23.8 | 36 |
| 7.4 | 7.1 | 12.3 | 12.8 | NA | 8 | 17.38 |
| 3.6 | 3.8 | 7.7 | 11.6 | NA | 5.2 | 18.19 |

The first column gives the known (mean) diameter values and the subsequent columns report the results from the automated analysis tools. Values are color-coded across their rows: values higher than the known (mean) diameter are depicted in shades of red (darker red means higher) and values lower than the known (mean) diameter are depicted in shades of green (darker green means lower). NA, not available.

**Supplementary Table VII**. Mean and median of fiber length per surface are of the image and the correlations between different tools in SEM images

| | Mean fiber length ± SD (µm/µm²) | Median [25th-75th percentile] (µm/µm²) | DiameterJ | AngioTool | REAVER | ER network analysis |
|---|---|---|---|---|---|---|
| **DiameterJ** | 2.60 ± 0.23 | 2.65 [2.51-2.77] | | 0.92* | 0.71* | 0.70* |
| **AngioTool** | 4.36 ± 0.30 | 4.36 [4.16-4.62] | 0.92* | | 0.76* | 0.69* |
| **REAVER** | 2.71 ± 0.25 | 2.77 [2.56-2.90] | 0.71* | 0.76* | | 0.64* |
| **ER network analysis** | 5.50 ± 0.54 | 5.48 [5.21-5.85] | 0.70* | 0.69* | 0.64* | |

Pearson correlation coefficient; *p<0.05. Green shading indicates significant association and no shading (white) indicates no significant association.

**Supplementary Table VIII**. Mean and median of fiber length per surface area of the image and the correlations between different tools in STED images

| | Mean fiber length ± SD (µm/µm²) | Median [25th-75th percentile] (µm/µm²) | DiameterJ | CT-FIRE | AngioTool | REAVER | ACCMetrics | ER network analysis | SOAX |
|---|---|---|---|---|---|---|---|---|---|
| **DiameterJ** | 0.21 ± 0.10 | 0.18 [0.14-0.23] | | 0.72* | 0.85* | 0.70* | 0.85* | 0.75* | 0.76* |
| **CT-FIRE** | 0.20 ± 0.08 | 0.19 [0.14-0.24] | 0.72* | | 0.65* | 0.83* | 0.56* | 0.48* | 0.50* |
| **AngioTool** | 0.40 ± 0.17 | 0.37 [0.28-0.48] | 0.85* | 0.65* | | 0.72* | 0.68* | 0.21 | 0.64* |
| **REAVER** | 0.22 ± 0.12 | 0.21 [0.15-0.28] | 0.70* | 0.83* | 0.72* | | 0.49* | 0.57* | 0.54* |
| **ACCMetrics** | 0.30 ± 0.11 | 0.28 [0.21-0.34] | 0.85* | 0.56* | 0.68* | 0.49* | | 0.59* | 0.95* |
| **ER network analysis** | 0.52 ± 0.48 | 0.37 [0.20-0.64] | 0.75* | 0.48* | 0.21 | 0.57* | 0.59* | | 0.57* |
| **SOAX** | 0.43 ± 0.14 | 0.42 [0.32-0.51] | 0.76* | 0.50* | 0.64* | 0.54* | 0.95* | 0.57* | |

Pearson correlation coefficient; *p<0.05. Green shading indicates significant association and no shading (white) indicates no significant association.

**Supplementary Table IX**. Mean and median of tools that quantify fiber length per surface area of the image and the correlations between different tools in confocal images

| | Mean fiber length ± SD (µm/µm²) | Median [25th-75th percentile] (µm/µm²) | DiameterJ | CT-FIRE | REAVER | ACCMetrics | ER network analysis |
|---|---|---|---|---|---|---|---|
| **DiameterJ** | 0.43 ± 0.04 | 0.43 [0.41-0.46] | | 0.84* | -0.01 | -0.22 | 0.89* |
| **CT-FIRE** | 0.83 ± 0.04 | 0.85 [0.82-0.96] | 0.84* | | 0.13 | -0.16 | 0.69* |
| **REAVER** | 0.17 ± 0.11 | 0.15 [0.07-0.26] | -0.01 | 0.13 | | 0.23 | 0.02 |
| **ACCMetrics** | 0.36 ± 0.03 | 0.36 [0.34-0.38] | -0.22 | -0.16 | 0.23 | | -0.36* |
| **ER network analysis** | 0.95 ± 0.20 | 0.97 [0.81-1.07] | 0.89* | 0.69* | 0.02 | -0.36* | |

Pearson correlation coefficient; *p<0.05. Green shading indicates significant association, orange shading indicates significant association in the opposite direction of what we expected, and no shading (white) indicates no significant association.

**Supplementary Table X**. Quantification of total fiber length in simulated networks

| Total fiber length (pixels) | DiameterJ | ER network analysis | AngioTool | REAVER |
|---|---|---|---|---|
| 94620 | 48537 | 85116 | 78425 | 56686 |
| 141720 | 91506 | 128565 | 92804 | 54703 |
| 178600 | 128297 | 165268 | 93807 | 35252 |

The first column gives the known total fiber length values and the subsequent columns report the results from the automated analysis tools. Values are color-coded: values lower than the known total fiber length are depicted in shades of green (darker green means lower).

**Supplementary Table XI**. Mean and median of the number of branch points per surface area of the image and the correlations between different tools in SEM images

| | Mean number of branch points ± SD (/μm²) | Median [25th-75th percentile] (/μm²) | DiameterJ | StructuralGT | AngioTool | REAVER | ER network analysis |
|---|---|---|---|---|---|---|---|
| **DiameterJ** | 8.62 ± 1.64 | 8.97 [7.87-9.68] | | 0.88* | 0.89* | 0.63* | 0.45* |
| **StructuralGT** | 5.82 ± 1.45 | 5.90 [4.68-7.06] | 0.88* | | 0.96* | 0.70* | 0.51* |
| **AngioTool** | 7.68 ± 1.44 | 7.81 [6.77-8.90] | 0.89* | 0.96* | | 0.71* | 0.46* |
| **REAVER** | 4.13± 0.87 | 4.36 [3.65-4.85] | 0.63* | 0.70* | 0.71* | | 0.46* |
| **ER network analysis** | 19.45 ± 3.29 | 19.36 [17.51-21.28] | 0.45* | 0.51* | 0.46* | 0.46* | |

Pearson correlation coefficient; *p<0.05. Green shading indicates significant association and no shading (white) indicates no significant association.

**Supplementary Table XII**. Mean and median of the number of branch points per surface area of the image and the correlations between different tools in STED images

| | Mean number of branch points ± SD (/μm²) | Median [25th-75th percentile] (/μm²) | DiameterJ | StructuralGT | AngioTool | REAVER | ACCMetrics |
|---|---|---|---|---|---|---|---|
| **DiameterJ** | 0.10 ± 0.09 | 0.07 [0.06-0.11] | | 0.87* | 0.89* | 0.72* | 0.77* |
| **StructuralGT** | 0.04 ± 0.03 | 0.04 [0.02-0.05] | 0.87* | | 0.85* | 0.64* | 0.86* |
| **AngioTool** | 0.09 ± 0.07 | 0.07 [0.05-0.11] | 0.89* | 0.85* | | 0.74* | 0.76* |
| **REAVER** | 0.06 ± 0.05 | 0.04 [0.03-0.08] | 0.72* | 0.64* | 0.74* | | 0.48* |
| **ACCMetrics** | 0.04 ± 0.02 | 0.04 [0.03-0.05] | 0.77* | 0.86* | 0.76* | 0.48* | |

Pearson correlation coefficient; *p<0.05. Green shading indicates significant association and no shading (white) indicates no significant association.

**Supplementary Table XIII**. Mean and median of the number of branch points per surface area of the image and the correlations between different tools in confocal images

| | Mean number of branch points ± SD (/µm$^2$) | Median [25th-75th percentile] (/µm$^2$) | DiameterJ | StructuralGT | REAVER | ACCMetrics | ER network analysis |
|---|---|---|---|---|---|---|---|
| **DiameterJ** | 0.34 ± 0.06 | 0.35 [0.30-0.39] | | 0.98* | 0.04 | -0.56* | 0.90* |
| **StructuralGT** | 0.14 ± 0.01 | 0.14 [0.13-0.15] | 0.98* | | 0.12 | -0.61* | 0.88* |
| **REAVER** | 0.03 ± 0.02 | 0.02 [0.01-0.05] | 0.04 | 0.12 | | 0.04 | -0.07 |
| **ACCMetrics** | 0.03 ± 0.01 | 0.03 [0.02-0.04] | -0.56* | -0.61* | 0.04 | | -0.60* |
| **ER network analysis** | 0.78 ± 0.26 | 0.85 [0.52-0.98] | 0.90* | 0.88* | -0.07 | -0.60* | |

Pearson correlation coefficient; *p<0.05. Green shading indicates significant association, orange shading indicates significant association in the opposite direction of what we expected, and no shading (white) indicates no significant association.

**Supplementary Table XIV**. Quantification of the number of branch points in synthetic images with known numbers

| Number of branch points | DiameterJ | ER network analysis | AngioTool | StructuralGT | REAVER |
|---|---|---|---|---|---|
| 745 | 992 | 1200 | 796 | 690 | 500 |
| 1654 | 2644 | 2501 | 1012 | 1357 | 520 |
| 2531 | 4469 | 3901 | 837 | 1755 | 283 |

The first column gives the known (mean) diameter values and the subsequent columns report the results from the automated analysis tools. Values are color-coded: values higher than the known (mean) diameter are depicted in shades of red (darker red means higher) and values lower than the known (mean) diameter are depicted in shades of green (darker green means lower).

**Supplementary Table XV**. Mean and median of the fibrin network density in SEM, STED, and confocal images

| | SEM | | STED | | Confocal | |
|---|---|---|---|---|---|---|
| | **Mean density ± SD (%)** | **Median [25th-75th percentile] (%)** | **Mean density ± SD (%)** | **Median [25th-75th percentile] (%)** | **Mean density ± SD (%)** | **Median [25th-75th percentile] (%)** |
| **AngioTool** | 50.3 ± 1.4 | 50.5 [49.3-51.3] | 10.8 ± 3.9 | 10.4 [8.1-12.8] | - | - |
| **REAVER** | 45.2 ± 6.8 | 44.9 [41.0-48.3] | 9.9 ± 6.0 | 8.2 [5.9-13.3] | 21.6 ± 14.5 | 18.4 [8.4-34.3] |
| **ACCMetrics** | - | - | 3.5 ± 1.3 | 3.2 [2.7-4.2] | 45.1 ± 4.5 | 46.3 [42.7-48.2] |

**Supplementary Table XVI**. Mean and median of the fractal dimension of the fibrin network in SEM, STED, and confocal images

| | SEM | | STED | | Confocal | |
|---|---|---|---|---|---|---|
| | **Mean fractal dimension ± SD** | **Median [25th-75th percentile]** | **Mean fractal dimension ± SD** | **Median [25th-75th percentile]** | **Mean fractal dimension ± SD** | **Median [25th-75th percentile]** |
| **BoneJ** | 1.80 ± 0.02 | 1.81 [1.79-1.81] | 1.37 ± 0.12 | 1.35 [1.28-1.42] | 1.88 ± 0.02 | 1.89 [1.87-1.90] |
| **Algorithms Hood** | 1.68 ± 0.02 | 1.68 [1.67-1.69] | 1.23 ± 0.05 | 1.22 [1.19-1.25] | 1.72 ± 0.06 | 1.74 [1.71-1.75] |
| **ACCMetrics** | - | - | 1.47 ± 0.09 | 1.49 [1.44-1.52] | 1.67 ± 0.02 | 1.67 [1.66-1.68] |

**Supplementary Table XVII**. Mean and median of the pore size or porosity and the correlations between different tools in confocal images

| | Mean ± SD | Median [25th-75th percentile] | Pore size analysis Krauss et al. (diameter) | Bubble analysis (diameter) | Hydrogel analysis (diameter) | Hydrogel analysis (area) | DiameterJ (area) | Hydrogel analysis (porosity) | Algorithms Hood et al. (porosity) | Quanfima (porosity) | DiameterJ (porosity) |
|---|---|---|---|---|---|---|---|---|---|---|---|
| **Pore size analysis Krauss et al. (diameter)** | 8.69 ± 1.53 | 8.40 [7.78-9.25] | | 0.09 | 0.36* | 0.39* | 0.15 | 0.07 | -0.03 | -0.13 | -0.02 |
| **Bubble analysis (diameter)** | 0.54 ± 0.08 | 0.55 [0.47-0.59] | 0.09 | | -0.14 | -0.13 | -0.08 | 0.09 | -0.07 | -0.00 | -0.03 |
| **Hydrogel analysis (diameter)** | 0.99 ± 0.11 | 0.98 [0.91-1.07] | 0.36* | -0.14 | | 0.98* | 0.49* | 0.45* | 0.38* | 0.60* | 0.62* |
| **Hydrogel analysis (area)** | 1.38 ± 0.37 | 1.32 [1.05-1.68] | 0.39* | -0.13 | 0.98* | | 0.56* | 0.42* | 0.45* | 0.64* | 0.67* |
| **DiameterJ (area)** | 3.74 ± 1.83 | 3.12 [2.70-4.16] | 0.15 | -0.08 | 0.49* | 0.56* | | -0.23 | 0.68* | 0.63* | 0.71* |
| **Hydrogel analysis (porosity)** | 0.48 ± 0.03 | 0.48 [0.47-0.50] | 0.07 | 0.09 | 0.45* | 0.42* | -0.23 | | -0.28 | 0.09 | -0.01 |
| **Algorithms Hood et al. (porosity)** | 0.67 ± 0.04 | 0.66 [0.65-0.68] | -0.03 | -0.07 | 0.38* | 0.45* | 0.68* | -0.28 | | 0.88* | 0.91* |
| **Quanfima (porosity)** | 0.62 ± 0.05 | 0.62 [0.58-0.64] | -0.13 | -0.00 | 0.60* | 0.64* | 0.63* | 0.09 | 0.88* | | 0.98* |
| **DiameterJ (porosity)** | 0.54 ± 0.03 | 0.54 [0.52-0.56] | -0.02 | -0.03 | 0.62* | 0.67* | 0.71* | -0.01 | 0.91* | 0.98* | |

Pearson correlation coefficient; *p<0.05. Green shading indicates significant association and no shading (white) indicates no significant association.

**Supplementary Table XVIII.** Preprocessing steps and settings for each tool

| Tool | SEM | STED | Confocal |
|---|---|---|---|
| **DiameterJ** | **(B)** T2 segmentation | **(B)** S2 segmentation | **(B) – (M)** M6 segmentation for whole image, T1 segmentation for 512x512 image (pore area) |
| **Local Thickness** | **(B)** | **(B)** | **x** |
| **SIMPoly** | No parameters adjusted | **x** | **x** |
| **AngioTool** | Threshold 15-255; vessel thickness 7; small particles 0; fill holes 0 | Threshold 115-255; vessel thickness 5; small particles 237; fill holes 0 | **x** |
| **StructuralGT** | Otsu Threshold; Blurring kernel size 5; Apply Gaussian Blur; Merge nearby nodes; Prune dangling edges; Assign edge weights by diameters; Remove disconnected segments; Remove object size 250; Disable Multigraph | **(B)** OTSU threshold; Blurring kernel size 21; Apply Gaussian blur; Merge nearby nodes; Remove disconnected segments; Remove object size 100 | **(B) – (M) A**daptive Threshold; Local threshold kernel 401; Blurring kernel size 3; Apply Gaussian blur; Merge nearby nodes; Prune dangling edges; Remove disconnected segments; Remove object size 500 |
| **REAVER** | **(B)** Grey threshold 0.21 or 0.29; Averaging Filter size 400 | Grey Threshold 0.02; Averaging Filter size 35 | **(M)** Grey Threshold 0.2; Averaging Filter size 30 |
| **BoneJ** | Binary DiameterJ images used, startboxsize 512, smallesboxsize 6, scalefactor 1.2, translations 0 | Binary DiameterJ images used, startboxsize 512, smallesboxsize 6, scalefactor 1.2, translations 0 | **(M)** Binary DiameterJ images used, startboxsize 512, smallesboxsize 6, scalefactor 1.2, translations 0 |
| **Algorithms Hood et al.** | **(B)** for fractal dimension: changed imfill →'holes' to 4 and numBlocks=1:25 to numBlocks=1:250 | for fractal dimension: changed numBlocks=1:25 to numBlocks=1:250 | **(M)** for fractal dimension: changed imfill →'holes' to 4 and numBlocks=1:25 to numBlocks=1:250 |
| **Bubble analysis** | **(B)** | **x** | **(M)** 512x512 image used, left upper corner of the image |
| **Quanfima** | **(B)** | **(B)** | **(B) – (M)** |
| **ER network analysis** | 0.0083 um/pix - process - template - Boundary T2 - Cisterna - Enhance - hyst +hmin Skeleton (0.3 - 0.5) – turn off Cisterna - Skeleton - Network extract change to distance - Width - Network - Save data | 0.026 um/pix – process – template - Boundary T2 – Cisterna – Enhance - Hysteresis skeleton - turn cisternae off – Skeleton - Network extract change to distance - Width - Network - Save data | **(B) – (M)** 0.06761 um/pix - process - template - Boundary T1 - Cisterna - Enhance - hyst +hmin Skeleton (0.3 - 0.6) - turn cisternae off – Skeleton - Network extract change to distance - Width - Network - Save data |

| | | | |
|---|---|---|---|
| **CT-FIRE** | **x** | thresh_im2=5; s_xlinkbox=8, fraction of coefs to keep=0.01; min fiber length=114 pixels (3000 nm); max fiber width=38 pixels (1000 nm) | **(M)** thresh_im2=5; s_xlinkbox=8, fraction of coefs to keep=0.01; min fiber length=30 pixels (2028 nm); max fiber width=15 pixels (1014 nm) |
| **ACCMetrics** | **x** | **(B)** 0.7 µm/pixel; images need to be a square | **(B) – (M)** 0.7 µm/pixel; images need to be a square |
| **Pore size analysis (Krauss et al.)** | **x** | **x** | 10.000 random points (512x512 image used) |
| **Hydrogel pore size analysis** | **x** | **x** | **(B) – (M)** |
| **SOAX** | **x** | **(B)** Intensity Scaling 0; Gaussian SD 2 pixels; Ridge threshold 0.02; minimum foreground 50; snake point spacing 1 pixel; minimum snake length 30 pixels | **x** |
| **Tool** | | | **Flow confocal** |
| **OrientationJ** | **x** | **x** | Local intensity derivatives in x and y were determined using a Gaussian window of 3 pixels and a Gaussian gradient. Calculation of nematic order parameter based on orientation measurements (Ψ): $S = \pm\sqrt{\langle\cos 2\psi\rangle^2} + \sqrt{\langle\sin 2\psi\rangle^2}$ |
| **Directionality plugin** | **x** | **x** | Local gradient mode |
| **CurveAlign** | **x** | **x** | Fiber analysis method: CT; Boundary method: No Boundary; fraction of coefs to keep = 0.01 |
| **FibLab / FOA tool** | **x** | **x** | 1 gaussian peak |
| **Cytospectre** | **x** | **x** | Component = mixed |
| **GTFiber** | **x** | **x** | Gaussian smoothing 387.68nm; orientation smoothing 1163.04nm; diffusion time 5s; top hat size 2000nm; Adaptive Threshold surface; Fringe |

| | | | |
|---|---|---|---|
| | | | Removal 1000nm; Grid step 10.000nm; Frame step 10.000nm |
| **FiberFit** | **x** | **x** | Low 68; High 512; Radial step 0.5; Angle increment 1 |
| **CT-FIRE** | **x** | **x** | **(B)** |
| **AFT (Alignment by Fourier Transform)** | **x** | **x** | Window size 100 pixels, overlap 50%; neighborhood radius 2; Filter blanks value 5 |

**(B)** means bandpass filter with 40 and 2 was used before using the image in the automated tool. **(M)** means a maximum projection was made before using the Z-stack in the automated tool. **x** indicates the tool was not used on the images from that column.

### Supplementary References


1. Vos BE, Martinez-Torres C, Burla F, Weisel JW, Koenderink GH. Revealing the molecular origins of fibrin's elastomeric properties by in situ X-ray scattering. *Acta Biomater*. 2020;104:39-52.
2. Ouyang W, Aristov A, Lelek M, Hao X, Zimmer C. Deep learning massively accelerates super-resolution localization microscopy. *Nat Biotechnol*. 2018;36(5):460-468.
3. Rigort A, Günther D, Hegerl R, et al. Automated segmentation of electron tomograms for a quantitative description of actin filament networks. *J Struct Biol*. 2012;177(1):135-144.
4. Usov I, Mezzenga R. FiberApp: An Open-Source Software for Tracking and Analyzing Polymers, Filaments, Biomacromolecules, and Fibrous Objects. *Macromolecules*. 2015;48(5):1269-1280.
5. Weissenböck J, Amirkhanov A, Li W, et al. FiberScout: An Interactive Tool for Exploring and Analyzing Fiber Reinforced Polymers. *2014 IEEE Pacific Visualization Symposium*; 2014:153-160.
6. Immink JN, Maris JJE, Capellmann RF, Egelhaaf SU, Schurtenberger P, Stenhammar J. ArGSLab: a tool for analyzing experimental or simulated particle networks. *Soft matter*. 2021;17(36):8354-8362.
7. Berg S, Kutra D, Kroeger T, et al. Ilastik: interactive machine learning for (bio)image analysis. *Nat Methods*. 2019;16(12):1226-1232.
8. Sympatec GmbH. PAQXOS; 2021.
9. Boudaoud A, Burian A, Borowska-Wykręt D, et al. FibrilTool, an ImageJ plug-in to quantify fibrillar structures in raw microscopy images. *Nat Protoc*. 2014;9(2):457-463.
10. Carpentier G, Berndt S, Ferratge S, et al. Angiogenesis Analyzer for ImageJ - A comparative morphometric analysis of "Endothelial Tube Formation Assay" and "Fibrin Bead Assay". *Sci Rep*. 2020;10(1):11568.
11. Sokolov PA, Belousov MV, Bondarev SA, Zhouravleva GA, Kasyanenko NA. FibrilJ: ImageJ plugin for fibrils' diameter and persistence length determination. *Comput Phys Commun*. 2017;214:199-206.
12. Bloksgaard M, Thorsted B, Brewer JR, De Mey JGR. Assessing Collagen and Elastin Pressure-dependent Microarchitectures in Live, Human Resistance Arteries by Label-free Fluorescence Microscopy. *J Vis Exp*. 2018(134):e57451.
13. Media Cybernetics. Fiber Thickness App; 2021.
14. Meijering E, Jacob M, Sarria JC, Steiner P, Hirling H, Unser M. Design and validation of a tool for neurite tracing and analysis in fluorescence microscopy images. *Cytometry A*. 2004;58(2):167-176.
15. Breuer D, Nowak J, Ivakov A, Somssich M, Persson S, Nikoloski Z. System-wide organization of actin cytoskeleton determines organelle transport in hypocotyl plant cells. *Proc Natl Acad Sci U S A*. 2017;114(28):E5741-E5749.
16. Kapoor V, Hirst WG, Hentschel C, Preibisch S, Reber S. MTrack: Automated Detection, Tracking, and Analysis of Dynamic Microtubules. *Sci Rep*. 2019;9(1):3794.
17. Smith MB, Li HS, Shen TA, Huang XL, Yusuf E, Vavylonis D. Segmentation and Tracking of Cytoskeletal Filaments Using Open Active Contours. *Cytoskeleton*. 2010;67(11):693-705.
18. Thermo Fisher Scientific. Phenom FiberMetric Software; 2021.
19. Wicker BK, Hutchens HP, Wu Q, Yeh AT, Humphrey JD. Normal basilar artery structure and biaxial mechanical behaviour. *Comput Methods Biomech Biomed Engin*. 2008;11(5):539-551.
20. Reis LA, Garcia APV, Gomes EFA, et al. Canine mammary cancer diagnosis from quantitative properties of nonlinear optical images. *Biomed Opt Express*. 2020;11(11):6413-6427.
21. Stevens CR, Berenson J, Sledziona M, Moore TP, Dong L, Cheetham J. Approach for semi-automated measurement of fiber diameter in murine and canine skeletal muscle. *PLoS ONE*. 2020;15(12):e0243163.
22. Kittisopikul M, Vahabikashi A, Shimi T, Goldman RD, Jaqaman K. Adaptive multiorientation resolution analysis of complex filamentous network images. *Bioinformatics.* 2020;36(20):5093-5103.

23. Morais FP, Carta A, Amaral ME, Curto JMR. Experimental 3D fibre data for tissue papers applications. *Data Brief*. 2020;30:105479.
24. Liao M, Liang X, Howard J. The narrowing of dendrite branches across nodes follows a well-defined scaling law. *Proc Natl Acad Sci U S A*. 2021;118(27).
25. Jiřík M, Tonar Z, Králíčková A, et al. Stereological quantification of microvessels using semiautomated evaluation of X-ray microtomography of hepatic vascular corrosion casts. *Int J Comput Assisted Radiol Surg*. 2016;11(10):1803-1819.
26. Ho SY, Chao CY, Huang HL, Chiu TW, Charoenkwan P, Hwang E. NeurphologyJ: An automatic neuronal morphology quantification method and its application in pharmacological discovery. *BMC Bioinform*. 2011;12.
27. Liu Y, Kolagunda A, Treible W, Nedo A, Caplan J, Kambhamettu C. Intersection To Overpass: Instance Segmentation On Filamentous Structures With An Orientation-Aware Neural Network And Terminus Pairing Algorithm. *IEEE Comput Soc Conf Comput Vis Pattern Recognit workshops*. 2019;2019:125-133.
28. Stanger JJ, Tucker N, Buunk N, Truong YB. A comparison of automated and manual techniques for measurement of electrospun fibre diameter. *Polym Test*. 2014;40:4-12.
29. Ovesný M, Křížek P, Borkovec J, Svindrych Z, Hagen GM. ThunderSTORM: a comprehensive ImageJ plug-in for PALM and STORM data analysis and super-resolution imaging. *Bioinformatics*. 2014;30(16):2389-2390.
30. Rossen NS, Kyrsting A, Giaccia AJ, Erler JT, Oddershede LB. Fiber finding algorithm using stepwise tracing to identify biopolymer fibers in noisy 3D images. *Biophys J*. 2021;120(18):3860-3868.
31. Zong Y, Pruner I, Antovic A, et al. Phosphatidylserine positive microparticles improve hemostasis in in-vitro hemophilia A plasma models. *Sci Rep*. 2020;10(1):7871.
32. Ghesquière P, Elsherbiny A, Fortier E, et al. An open-source algorithm for rapid unbiased determination of DNA fiber length. *DNA Repair*. 2019;74:26-37.
33. Faulkner C, Zhou J, Evrard A, et al. An automated quantitative image analysis tool for the identification of microtubule patterns in plants. *Traffic*. 2017;18(10):683-693.
34. Zhang RM, Kumra H, Reinhardt DP. Quantification of Extracellular Matrix Fiber Systems Related to ADAMTS Proteins. *Methods Mol Biol*. Vol. 2043; 2020:237-250.
35. Shah Hosseini N, Simon B, Messaoud T, Khenoussi N, Schacher L, Adolphe D. Quantitative approaches of nanofibers organization for biomedical patterned nanofibrous scaffold by image analysis. *J Biomed Mater Res Part A*. 2018;106(11):2963-2972.
36. Seet LF, Chu SWL, Teng X, Toh LZ, Wong TT. Assessment of progressive alterations in collagen organization in the postoperative conjunctiva by multiphoton microscopy. *Biomed Opt Express*. 2020;11(11):6495-6515.
37. Canver AC, Morss Clyne A. Quantification of Multicellular Organization, Junction Integrity, and Substrate Features in Collective Cell Migration. *Microsc Microanal*. 2017;23(1):22-33.
38. Xu T, Langouras C, Koudehi MA, et al. Automated Tracking of Biopolymer Growth and Network Deformation with TSOAX. *Sci Rep*. 2019;9(1):1717.
39. Zhang Z, Xia S, Kanchanawong P. An integrated enhancement and reconstruction strategy for the quantitative extraction of actin stress fibers from fluorescence micrographs. *BMC Bioinform*. 2017;18(1):268.
40. Eekhoff JD, Lake SP. Three-dimensional computation of fibre orientation, diameter and branching in segmented image stacks of fibrous networks. *J R Soc Interface*. 2020;17(169):20200371.
41. Zhang Z, Nishimura Y, Kanchanawong P. Extracting microtubule networks from superresolution single-molecule localization microscopy data. *Mol Biol Cell*. 2017;28(2):333-345.
42. Flormann DAD, Schu M, Terriac E, et al. A novel universal algorithm for filament network tracing and cytoskeleton analysis. *FASEB J*. 2021;35(5):e21582.
43. Windoffer R, Schwarz N, Yoon S, et al. Quantitative Mapping of Keratin Networks in 3D. *eLife*. 2022;11.

44. Wisanpitayakorn P, Mickolajczyk KJ, Hancock WO, Vidali L, Tüzel E. Measurement of the Persistence Length of Cytoskeletal Filaments using Curvature Distributions. *Biophys J*. 2022:121(10):1813-1822.
45. Lee T, Barone T, Rubinstein E, Mischler S. Asbestos fiber length and width comparison between manual and semi-automated measurements. *J Occup Environ Hyg*. 2022:1-12.
46. Wershof E, Park D, Barry DJ, et al. A FIJI macro for quantifying pattern in extracellular matrix. *Life Sci Alliance*. 2021;4(3).
47. Du P, Ding X, Liang Q, Chen X, Zhang Y, Chen L. Numerical and experimental analysis on anisotropic thermal conductivity of carding quartz fiber web based on the reconstruction model. *Text Res J*. 2022;92(11-12):2100-2111.

### Literature search terms

(collagen/exp OR 'collagen gel'/de OR fibrin/exp OR fiber/de OR 'collagen fiber'/de OR microtubule/de OR 'molecular scaffold'/de OR (collagen* OR fibrin* OR (((fiber* OR fibre*) NEAR/3 (orientation* OR alignment* OR diameter* OR length OR organization* OR organisation* OR extraction* OR radius* OR polymer*)) NOT nerve-fib*) OR nanofiber* OR nanofibr* OR microfiber* OR microfibr* OR microtubul* OR nanotubul* OR tubules OR (network* NEAR/3 (orientation* OR quantif*)) OR ((fibrillar OR tubular OR polymer* OR fibroid* OR nanofibr* OR microfibr* OR hydrogel* OR biomaterial* OR fibre* OR fiber* OR composite*) NEAR/3 scaffold*) OR nanotube* OR microtube*):ab,ti)

AND

('quantitative analysis'/de OR orientation/de OR anisotropy/de OR thickness/de OR 'structure analysis'/de OR ultrastructure/de OR (orientation* OR alignment* OR diameter* OR network* OR quantificat* OR quantitat* OR topolog* OR architecture* OR anisotrop* OR thickness* OR ((fiber* OR fibre*) NEAR/3 (length OR organization* OR organisation* OR extraction* OR radius* OR branchpoint* OR branch-point* OR Intersection* OR Inter-section* OR connectivit*)) OR (physical* NEAR/3 propert*) OR ((structur* OR microstructur*) NEAR/3 analy*) OR ultrastructure* OR pore-size OR porosit*):Ab,ti)

AND

('image analysis'/de OR automation/de OR software/de OR algorithm/de OR 'mathematical analysis'/de OR 'mathematical computing'/de OR 'computer analysis'/de OR 'data analysis software'/de OR autoanalysis/de OR 'automated pattern recognition'/de OR ((image* NEAR/3 (analys*)) OR automat* OR software* OR soft-ware* OR algorithm* OR (mathematic* NEAR/3 (analysis OR computing)) OR (computer* NEAR/3 analy*) OR autoanaly* OR (pattern* NEAR/3 recognition*)):ab,ti)

AND

(microscopy/exp OR fluorescence/exp OR 'fluorescence imaging'/exp OR 'fluorescence imaging system'/exp OR Polarimetry/de OR  polarization/de OR 'three-dimensional imaging'/de OR refractometry/de OR refractometer/de OR (microscop* OR polarization* OR polarisation* OR polarized-light* OR polarised-light* OR polarimetr* OR fluorescen* OR three-dimensional* OR 3d OR 3-d OR refractomet*):Ab,ti)

NOT

[conference abstract]/lim

AND

[english]/lim

#### Python code for simulated fibrous networks

```
##########################################################################
# Simulation of fibrous networks on a 2D surface                          #
# Felix Frey, TU Delft, 2022                                              #
##########################################################################

#Algorithm to generate fibrous networks:
#For every network simulation, we set six parameters:
#(1) the total number of filaments “tot_number_fil”,
#(2) the maximum filament length “max_len”,
#(3) and (4) two parameters “shape” and “scale” that define the filament
#diameter (explained below), and
#(5) and (6) two parameters “geometry_x” and “geometry_y” that determine
#the length and height of the simulated surface,
#which are both set to 100 for all simulated networks.

#We then distribute the total number of filaments on the surface.
#First, we determine the filament orientation. For every filament, we draw a
#random angle ϕ from a uniform distribution between 0 and π with respect to
#the x-axis. Second, we determine the position of the filament center of mass.
#Therefore, for every filament, we draw the x and y coordinate of the
#filament center of mass from a uniform distribution between 0 and
#“geometry_x” and between 0 and “geometry_y”, respectively. With the filament
#orientation and the position of the filament center of mass, we then
#calculate the length of the filament within the simulated surface.
#Finally, we set the filament diameter, by sampling a random number from a
#gamma distribution that is determined by the “shape” and “scale” parameters.

#After we have distributed the total number of filaments on the surface,
#we calculate the branch points for the network by summing up all
#intersections between pairs of two filaments within the simulated surface.
#From the simulation we determine the filament length and diameter
#distributions and the number of branch points.

#import used packages

import numpy as np
import matplotlib.pyplot as plt
from matplotlib.lines import Line2D
import warnings

#function that distributes a filament on the 2D surface.
#input: is the maximum filament length: max_len and the length and height
#of the simulated surface: geometry_x and geometry_y
#output: coordinates of the filament endpoints: x_0,y_0,x_1,y_1,
#the filament length: rod_length, the filament orientation: phi,
#the filament center of mass coordinates: x_nc and y_nc
#and the filament diameter: diameter

def DefineRod(max_len,geometry_x,geometry_y):

    rod_length=max_len
    #the filament orientation is drawn from a uniform distribution
    #between 0 and pi with respect to the x-axis.
    phi=np.random.rand()*np.pi

    #the filament center of mass position is drawn from a uniform distribution
    #between 0 and the length and height of the simulated surface
```

```
    x_nc=np.random.rand()*geometry_x
    y_nc=np.random.rand()*geometry_y

    #calculation of the filament coordinates within the simulated surface
    x_0,y_0,x_1,y_1=CaculateCoordinates(rod_length,phi,x_nc,y_nc)
    #filament length within the sumualte surface
    rod_length=np.sqrt((x_1-x_0)**2+(y_1-y_0)**2)

    #the filament diameter is drawn from a gamma distribution with shape
    #and scale parameter
    diameter=np.random.gamma(shape=shape,scale=scale)

    return x_0,y_0,x_1,y_1,rod_length,phi,x_nc,y_nc,diameter

#function that calculates the filament coordinates within the simulated surface
#input: filament length: length, the filament orientation: phi
#and the coordinates of the filament center of mass: x_nc and y_nc
#output: coordinates of the filament endpoints: x_0,y_0,x_1,y_1,

def CaculateCoordinates(length,phi,x_nc,y_nc):
    #define filament coordinates (includes space outside of the simulated surface)
    x_0=np.cos(phi)*length/2+x_nc
    y_0=np.sin(phi)*length/2+y_nc

    x_1=-np.cos(phi)*length/2+x_nc
    y_1=-np.sin(phi)*length/2+y_nc

    #orient the filament so that x_0 is the left coordinate and x_1 is the
    #right coordinate
    if x_0>x_1:
        x_temp=x_0
        x_0=x_1
        x_1=x_temp

        y_temp=y_0
        y_0=y_1
        y_1=y_temp

    #calcuate the filament slope m_g and the filament y-intercept c_g
    [m_g,c_g]=LineCalculator(x_0,y_0,x_1,y_1)

    #cacluate the filament coordinates within the simulated surface
    if x_0<0:
        x_0=0
        y_0=c_g
    if x_0>geometry_x:
        x_0=geometry_x
        y_0=geometry_x*m_g+c_g
    if y_0<0:
        y_0=0
        x_0=-c_g/m_g
    if y_0>geometry_y:
        y_0=geometry_y
        x_0=(geometry_y-c_g)/m_g

    if x_1<0:
        x_1=0
        y_1=c_g
    if x_1>geometry_x:
```

```
        x_1=geometry_x
        y_1=geometry_x*m_g+c_g
    if y_1<0:
        y_1=0
        x_1=-c_g/m_g
    if y_1>geometry_y:
        y_1=geometry_y
        x_1=(geometry_y-c_g)/m_g

    return x_0,y_0,x_1,y_1

#function that calculates the filament slope and the filament y-intercept
#input: coordinates of the filament endpoints: x_0,y_0,x_1,y_1
#output: the filament slope: m_g and y-intercept c_g

def LineCalculator(x_0,y_0,x_1,y_1):
    m_g=(y_1-y_0)/(x_1-x_0)
    c_g=y_1-m_g*x_1
    return [m_g,c_g]

#function that appends the filament properties to lists

def
UpdateStep(x_l,y_l,x_0,x_1,y_0,y_1,phi_l,phi,rod_len_l,rod_len,nc_l,x_nc,y_nc,diam
eter_l,diameter):

    x_l=np.append(x_l,np.array([x_0,x_1]))
    y_l=np.append(y_l,np.array([y_0,y_1]))
    phi_l=np.append(phi_l,np.array([phi]))
    rod_len_l=np.append(rod_len_l,np.array([rod_len]))
    nc_list=np.append(nc_l,np.array([x_nc,y_nc]))
    diameter_l=np.append(diameter_l,np.array([diameter]))
    return [x_l,y_l,phi_l,rod_len_l,nc_list,diameter_l]

#function that calculates the number of branching points=points of filament
#intersections of the simulated network
#input: lists of coordinates of the filament endpoints: pos_x_l,pos_y_l
#output: number of branching points

def BranchingPoints(pos_x_l,pos_y_l):
    counter=0
    #caculate for every filament the number of filament intersections within
    #the surface
    for i in range (tot_number_fil):

counter+=CheckIntersection(pos_x_l[2*i],pos_y_l[2*i],pos_x_l[2*i+1],pos_y_l[2*i+1]
,pos_x_l,pos_y_l)
    #since the intersection points are calculate twice for pairwise filament
    #intersection, we divide by 2.
    return int(counter/2)

#function that calculates for every filament the number of filament
#intersections within the surface
#input: coordinates of the filament endpoints: x_0,y_0,x_1,y_1
#and lists of coordinates of the filament endpoints: pos_x_l,pos_y_l
#output: number of branching points if any

def CheckIntersection(x_0,y_0,x_1,y_1,pos_x_list,pos_y_list):
    number_of_rods=int(len(pos_x_list)/2)
```

```
    if(number_of_rods)>0:

branching_points=np.sum(CalculateIntersction(x_0,y_0,x_1,y_1,pos_x_list,pos_y_list
))
        return branching_points
    else:
        return False

#function that first calculates all intersection points between the input
#filament and all other filaments of the network. Second, the function checks
#if the intersection points are part of the filaments within the surface
#the calculation is done in a parallel manner.
#input: coordinates of the filament endpoints: x_0,y_0,x_1,y_1
#and lists of coordinates of the filament endpoints: pos_x_l,pos_y_l
#output: an array of all True and False values indicating with which filaments
#the considered filament intersects

def CalculateIntersction(x_0,y_0,x_1,y_1,pos_x_list,pos_y_list):

    #two lines g: y_g=m_g*x+c_g and h: y_h=m_h*x+c_h intersect,
    #when x and y_g=y_h are identical -> x=(c_h-c_g)/(m_g-m_h)

    number_of_rods=int(len(pos_x_list)/2)
    #calculate the slope and y-intercept of the considered filament
    [m_g,c_g]=LineCalculator(x_0,y_0,x_1,y_1)

    #create arrays for the calculation
    m_g=np.full((1,number_of_rods), m_g)[0]
    c_g=np.full((1,number_of_rods), c_g)[0]
    x_0=np.full((1,number_of_rods), x_0)[0]
    x_1=np.full((1,number_of_rods), x_1)[0]

    #calculate the coordinates of the other filaments
    x0=pos_x_list[0::2]
    x1=pos_x_list[1::2]
    y0=pos_y_list[0::2]
    y1=pos_y_list[1::2]

    #calculate the slope and y-intercept of the other filaments
    #self-intersections (a filament with itself) will lead to a NaN value for
    #x_insec that are filtered out in the next step
    [m_h,c_h]=LineCalculator(x0,y0,x1,y1)
    x_insec=(c_h-c_g)/(m_g-m_h)
    y_insec=x_insec*m_g+c_g

    #check for intersections within the surface in parallel for all filaments
    #self-intersections are filtered out
    a=np.multiply(x_0 < x_insec,x_insec < x_1)
    aprime=np.multiply(0 < x_insec,x_insec < geometry_x)
    b=np.multiply(x0 < x_insec,x_insec < x1)
    bprime=np.multiply(0 < y_insec,y_insec < geometry_y)

    #return the intersection array that contains True and False values depending
    #on whether the considered filament intersects with the other filaments
    array_intersec=np.multiply(a*aprime, b*bprime)
    return array_intersec


#function that simulates the distribution of filaments on the 2D surface
```

```
#input: maximum filament length: max_len, the length and height
#of the simulated surface: geometry_x and geometry_y and the number of filaments:
#tot_number_fil
#output: arrays for the x- and y-coordinates of the endpoints of the filaments
#and the center of mass coordinates:
#pos_x_l, pos_y_l, nc_l
#and the orientation, length and diameter: phi_l, rod_len_l, diameter_l

def MikadoModel(max_len,geometry_x,geometry_y,tot_number_fil):
    pos_x_l,pos_y_l,phi_l,rod_len_l,nc_l,diameter_l=\

np.array([]),np.array([]),np.array([]),np.array([]),np.array([]),np.array([])
    number_fil=0

    #define filaments iteratively
    while number_fil<tot_number_fil:


x_0,y_0,x_1,y_1,rod_len,phi,x_nc,y_nc,diameter=DefineRod(max_len,geometry_x,geomet
ry_y)

        [pos_x_l,pos_y_l,phi_l,rod_len_l,nc_l,diameter_l]=\

UpdateStep(pos_x_l,pos_y_l,x_0,x_1,y_0,y_1,phi_l,phi,rod_len_l,rod_len,nc_l,x_nc,y
_nc,diameter_l,diameter)
        number_fil+=1

    return pos_x_l,pos_y_l,phi_l,rod_len_l,nc_l,diameter_l


#the class "LineDataUnits" was written by StackOverflow user
#ImportanceOfBeingErnest
(https://stackoverflow.com/users/4124317/importanceofbeingernest),
#published at https://stackoverflow.com/a/42972469
#under the license CC BY-SA 4.0 (https://creativecommons.org/licenses/by-sa/4.0/).
#the class allows to plot the sampled filament diameter in the same length unit
#as the filament length

class LineDataUnits(Line2D):
    def __init__(self, *args, **kwargs):
        _lw_data = kwargs.pop("linewidth", 1)
        super().__init__(*args, **kwargs)
        self._lw_data = _lw_data

    def _get_lw(self):
        if self.axes is not None:
            ppd = 72./self.axes.figure.dpi
            trans = self.axes.transData.transform
            return ((trans((1, self._lw_data))-trans((0, 0)))*ppd)[1]
        else:
            return 1

    def _set_lw(self, lw):
        self._lw_data = lw

    _linewidth = property(_get_lw, _set_lw)

#function that plots the filaments on the 2D surface
```

```
def PlotRods(filename,pos_x_list,pos_y_list,geometry_x,geometry_y,diameter_l):

    fig = plt.figure(figsize=(10,10))
    sub1=fig.add_subplot(111)

    for i in range(int(len(pos_x_list)/2)):


line=LineDataUnits([pos_x_list[i*2+0],pos_x_list[i*2+1]],[pos_y_list[i*2+0],pos_y_
list[i*2+1]],linewidth=diameter_l[i],color="r")
        sub1.add_line(line)

    sub1.set_aspect('equal')
    sub1.axis('off')
    sub1.set_xlim(0,geometry_x)
    sub1.set_ylim(0,geometry_y)
    plt.savefig("FibrousNetwork"+str(filename)+".pdf",bbox_inches='tight')

#function that plots the filament length distribution

def PlotLengthDistribtion(filename,rod_length_list,max_len):

    fig=plt.figure(figsize=(7*1.6,7))
    sub1=fig.add_subplot(111)


sub1.hist(rod_length_list,bins=int(np.sqrt(len(rod_length_list)))*2,alpha=0.5,face
color=new_colors[3])

    sub1.set_xlabel("length (length units)",fontsize=nn)
    sub1.set_ylabel("frequency",fontsize=nn)
    sub1.xaxis.set_tick_params(labelsize=nn)
    sub1.yaxis.set_tick_params(labelsize=nn)
    sub1.set_xlim(0,max_len*1.05)

    plt.tight_layout()
    plt.subplots_adjust(top=0.92)
    plt.savefig("LengthDistribution"+str(filename)+".pdf")

#function that plots the filament diameter distribution

def PlotWidthDistribtion(filename,diameter_length_list,max_dia):

    fig=plt.figure(figsize=(7*1.6,7))
    sub1=fig.add_subplot(111)


sub1.hist(diameter_length_list,bins=int(np.sqrt(len(diameter_length_list)))*2,alph
a=0.5,facecolor=new_colors[3])

    sub1.set_xlabel("diameter (length units)",fontsize=nn)
    sub1.set_ylabel("frequency",fontsize=nn)
    sub1.xaxis.set_tick_params(labelsize=nn)
    sub1.yaxis.set_tick_params(labelsize=nn)
    sub1.set_xlim(0,max_dia*1.05)


    plt.tight_layout()
    plt.subplots_adjust(top=0.92)
```

```
    plt.savefig("DiameterDistribution"+str(filename)+".pdf")

#function that saves the simulation parameters

def
WriteSimulationParameters(filename,max_len,tot_number_fil,shape,scale,branching_po
ints):
    np.savetxt("SimulationParameters"+filename+'.csv',
np.c_[max_len,tot_number_fil,shape,scale,branching_points],header="max_len,tot_num
ber_fil,shape,scale,branching_points", delimiter=',')


#plot settings

#colors
new_colors = ['#1f77b4', '#ff7f0e', '#2ca02c', '#d62728',
              '#9467bd', '#8c564b', '#e377c2', '#7f7f7f',
              '#bcbd22', '#17becf']
#fontsize
nn=26

########################################################################
#main function
########################################################################

#parameter values

tot_number_fil=100
geometry_x=100
geometry_y=100
max_len=geometry_x*np.sqrt(2)*2
shape=2
scale=0.1


#name of the data file
filename=str(3)

#function calls and definition of arrays for
#the filament coordinates: pos_x_list,pos_y_list
#the filament orientations and filament lengths: phi_list,rod_length_list
#the filament center of mass coordinates: nc_list
#and the filament diameter: diameter_list

#ignore runtime warnings related to self-intersections
warnings.filterwarnings("ignore", message="invalid value encountered in
true_divide")

#run main function
pos_x_list,pos_y_list,phi_list,rod_length_list,nc_list,diameter_list=MikadoModel(m
ax_len,geometry_x,geometry_y,tot_number_fil)
branching_points=BranchingPoints(pos_x_list,pos_y_list)
print("branching points: " + str(branching_points))
PlotRods(filename,pos_x_list,pos_y_list,geometry_x,geometry_y,diameter_list)
PlotLengthDistribtion(filename,rod_length_list,np.max(rod_length_list))
PlotWidthDistribtion(filename,diameter_list,np.max(diameter_list))
WriteSimulationParameters(filename,max_len,tot_number_fil,shape,scale,branching_po
ints)
```
